# Novel sensor-integrated proteome on chip (SPOC) platform with thousands of folded proteins on a 1.5 sq-cm biosensor chip to enable high-throughput real-time label-free screening for kinetic analysis

**DOI:** 10.1101/2024.01.23.575909

**Authors:** Chidozie Victor Agu, Rebecca L Cook, William Martelly, Lydia R Gushgari, Mukilan Mohan, Bharath Takulapalli

## Abstract

An automated proteomic platform for producing and screening an array of functional proteins on biosensor surfaces was developed to address the challenges of measuring proteomic interaction kinetics in high throughput (HTP). This technology is termed Sensor-Integrated Proteome On Chip (SPOC®) which involves *in-situ* cell-free protein expression in nano-liter volume wells (nanowells) directly from rapidly customizable arrays of plasmid DNA, facilitating simultaneous capture-purification of up to 2400 unique full-length folded proteins onto a 1.5 sq-cm surface of a single gold biosensor chip. Arrayed SPOC sensors can then be screened by real-time label-free analysis, including surface plasmon resonance (SPR) to generate kinetic affinity, avidity data. Fluorescent and SPR assays were used to demonstrate zero crosstalk between protein spots. The functionality of the SPOC protein array was validated by antibody binding assay, post-translational modification, mutation-mediated differential binding kinetics, and catalytic activity screening on model SPOC protein arrays containing p53, Src, Jun, Fos, HIST1H3A, and SARS-CoV-2 receptor binding domain (RBD) protein variants of interest, among others. Monoclonal antibodies were found to selectively bind their target proteins on the SPOC array. A commercial anti-RBD antibody was used to demonstrate discriminatory binding to numerous SARS-CoV-2 RBD variants of concern with comprehensive kinetic information. With advantages of HTP, flexibility, low-cost, quick turnaround time, and real-time kinetic affinity profiling, the SPOC proteomic platform addresses the challenges of interrogating protein interactions at scale and can be deployed in various research and clinical applications.

## Introduction

Protein interactions are core to innumerable biological functions that occur within the complex milieu of a cell and determine many core processes such as DNA transcription and translation, metabolic processing, regulation of enzyme turnover, intra-and inter-cellular signal transduction, and motor functions and cytoskeletal rearrangement, among many others. These protein interactions are characterized by unique binding affinities determined by a dynamic balance between association and dissociation rates amongst free and bound complexes. As a result, measuring the kinetic parameters of protein interactions with other proteins, DNA, RNA, metabolites and other biomolecules is critical to the functional characterization of biological interactions, development of more effective drug molecules and vaccines, and the discovery of new biomarkers and phenotypes. Therefore, to comprehensively characterize dysregulated protein interaction networks for disease diagnostics, and to assess the potency and safety of drug candidates during preclinical development, next-generation proteomic tools must be capable of capturing the kinetics of numerous and diverse protein interactions.

Functional protein characterization through measurement of interaction kinetics is key in many disease areas, including cancer, neurodegenerative diseases, and autoimmunity. In cancer, driver mutations can have a variety of effects on protein activity commonly leading to gain or loss of function, and acquisition of drug resistance. Across over 33 different cancer types and subtypes, missense mutations have been observed to be disproportionately enriched at protein-protein interfaces, indicating that mutations that disrupt interaction networks between proteins are common and would require kinetic screening to unravel their impacts in detail (Cheng et al. 2021). In this context, high-throughput proteomics technologies with the capability to characterize hundreds of cancer-associated mutations and isoforms at the level of protein function would hasten the mechanistic understanding of the consequences of these mutations, and further inform and improve the accuracy of advanced computational tools for driver mutation identification. In regards to the immune system, binding kinetics play a big role in evaluating important immune parameters such as infectivity of a new viral variant, efficacy of a new drug/neutralizing antibody, and immune parameters related to the development of autoimmunity such as the efficiency of central tolerance, efficiency of T-cell priming, and the efficiency of destroying a target cell expressing an autoantigen (Koehli et al., 2014). Neurodegenerative diseases, such as Alzheimer’s, Parkinson’s, and Huntington’s, are characterized by the abnormal accumulation and aggregation of specific proteins within the brain, leading to the formation of pathological structures. Understanding the kinetics of protein interactions, including their rates of synthesis, folding, misfolding, aggregation, and degradation, is crucial for unraveling the molecular mechanisms involved in neurodegenerative diseases. Kinetic studies enable researchers to elucidate the temporal aspects of these processes, helping to identify critical stages and potential therapeutic targets. In pharmacology and drug development, the kinetics of protein-drug interactions are crucial for determining the pharmacodynamics of therapeutic compounds. Likewise, in diagnostic applications, kinetic information can reveal additional information that current endpoint protein assays are unable to resolve, such as early detection and disease sub-typing. For example, in autoimmune disease diagnostics, autoantibodies (AAb) which have characteristic binding strength for their target antigens (avidities/affinities) may be predictive of disease pathogenesis and severity. For instance, in systemic autoimmune rheumatic diseases the presence of low affinity anti-citrullinated protein antibodies (ACPAs) among patients with rheumatoid arthritis is linked to increased presence of AAb synovial tissue penetration and more severe joint destruction (van der Woude et al., 2010; Suwannalai et al., 2014). This kind of information is unavailable via endpoint protein assays.

Drug screening and vaccine development, infectious disease surveillance and diagnostics efforts would benefit greatly from availability of technology platform for large-scale kinetic profiling of protein interactions. Unfortunately, despite the significant progress in proteomics, there remains a notable deficit in tools available for measuring protein binding kinetics in high throughput (HTP). Although fluorescence-based protein microarrays are commonly used for protein research, they are primarily suited for qualitative and semi-quantitative analysis where a relatively simple, low-medium-high scale of protein binding is needed. These do not output quantitative data needed for kinetic analysis. Protein interactions are transient equilibrium reactions and by leveraging label-free real-time biosensing approaches it becomes possible to directly measure kinetic parameter such as rates of association (*k*a) and dissociation (*k*d), dissociation half-life (t1/2), maximal feasible signal generated by the interaction between a ligand/analyte pair (Rmax), etc. The equilibrium dissociation constant (KD) or affinities, avidities can then be computed from these kinetic parameters, which can otherwise only be approximated by fluorescence-based approaches (Nagata et al., 2000). Various approaches exist for measuring protein kinetics, each with distinct advantages and limitations. Isothermal titration calorimetry (ITC) offers detailed thermodynamic information, such as binding constants, enthalpy, and entropy changes, but is low throughput and requires large sample input, making it less suitable for weak or fast-binding systems (Wang et al., 2020). Fluorescence Resonance Energy Transfer (FRET) allows for real-time monitoring of conformational changes and interactions, particularly in live cells, but relies on fluorescent labeling, which is prone to photobleaching and background fluorescence (Liao et al., 2020) and is not fully compatible with HTP kinetic screening. Stopped-Flow spectroscopy is adept at studying rapid reactions on a millisecond timescale but is limited to fast processes and may demand a substantial sample size (Zheng et al., 2015). Another method, Electrophoretic Mobility Shift Assay (EMSA) is a simple and cost-effective technique measuring changes in protein migration through a gel matrix in response to binding to specific ligands like nucleic acids. However, it primarily applies to specific interactions, such as DNA-protein binding, and may not provide kinetic data (Hellman and Fried, 2007). Nuclear Magnetic Resonance (NMR) Spectroscopy detects changes in nuclear magnetic resonance signals during protein-ligand interactions, offering structural, dynamic, and kinetic insights (Maity et al., 2019). However, NMR necessitates stable isotopic labeling, may not suit all systems, and is often constrained to relatively small proteins or complexes and is not an HTP method. Bio-layer interferometry (BLI) is another recent technique that monitors changes in interference patterns due to binding interactions between immobilized ligands and analyte proteins (Shah and Duncan, 2014). BLI is real-time and label-free and attractively available in high-throughput (HTP) instrument formats capable of monitoring interactions from 96-and 384-well plates, however the throughput can be at the cost of sensitivity and reproducibility compared to other, well-established label-free formats (Yang et al. 2016). Surface plasmon resonance (SPR) stands at the forefront of protein kinetics measurement, measuring changes in refractive index as molecules, such as proteins, bind and dissociate from a biomolecule or ligand coated gold biosensor surface. This method offers real-time, label-free, quantitative data, including kinetics and affinity information. However, this technique demands specialized equipment, which can be costly. Current state-of-the-art SPR systems, like GE’s Biacore, excel in quantitative protein binding analysis but has limitations in terms of throughput (Maity et al., 2019; Shah and Duncan, 2014).

The lack of accessible proteomic tools to produce an array of full-length proteins and measure binding kinetics in HTP hinders the ability to comprehensively study and understand the dynamics of protein interactions in complex biological systems. Importantly, the high cost of producing pure, fully folded and functional protein microarrays remains a challenge. Current methods for producing protein arrays rely on the laborious, time consuming and often cost-prohibitive method of expressing individual recombinant proteins in a cell-based or cell-free system, followed by purification and manual or robotic spotting of the affinity-purified recombinant proteins. Protein purification is time-consuming and expensive, and with purified commercial proteins costing upwards of $250 per protein, many labs lacking the proper infrastructure or expertise are locked out of functional proteomic studies. In terms of stability, traditional methods require low-temperature storage conditions to maintain the structural integrity and activity of recombinant proteins throughout the entire process, increasing costs and often leading to loss of materials and sample integrity. Consequently, cost can often add up significantly when scaled to screening thousands of proteins. Therefore, traditional methods of expression and robotic immobilization of individual proteins, either purified or from lysate, to produce functional protein arrays lack scalability, flexibility, versatility and customizability and cannot feasibly address challenges of measuring proteomic interaction kinetics in HTP. Promisingly, cell-free expression such as nucleic acid programmable protein array technology (NAPPA; Takulapalli et al., 2012; Ramachandran et al., 2004; Link and LaBaer et al., 2008a; Link and LaBaer et al., 2008b; Yu et al., 2017) and Isolated Protein Capture (IPC; Karthikeyan et al., 2016) represent significant milestones in protein microarray production methods that have considerably reduced the cost of protein microarray production while enabling facile array customization. However, measuring kinetic analysis of thousands of protein interactions simultaneously remains a challenge as most cell-free protein microarray platforms, including traditional NAPPA technology, are designed for label-based end-point detection methods such as fluorescence or luminescence. The reliance on these label-based end-point readout of analyte binding to arrayed proteins implies only a semi-quantitative and indirect measure of any given interaction.

Given the relevance of kinetics in proteomics it has become increasingly important to develop flexible, scalable proteomic tools with multiplex and HTP capabilities that combine scaled production of proteins, real-time label-free biosensing and kinetic analysis. One way to achieve this important capability is by integrating recent advances in SPR biosensing with *in situ* cell-free expression and IPC approaches for making and screening functional protein arrays. Recently, researchers combined cell-free protein expression and SPR to study protein interactions using mutants of p53 and oncoprotein MDM2 as a model system (Fuentes et al., 2021). Their method circumvents IPC by directly printing plasmid DNAs for expressing various p53 mutants on a gold SPR sensor as in traditional NAPPA on glass. Another SPR method involves transcription and translation of RNA from surface-bound double-stranded DNA (dsDNA) templates for the on-chip multiplexed biosynthesis of aptamer and protein microarrays in a microfluidic format (Fasoli and Corn, 2015). However, the notable caveat with these methods is that the presence of impurities (components of the print mix and DNA) can impair downstream kinetic analysis by SPR due to introduction of components that crowd the evanescent field wave which can reduce sensitivity, block access to the ligand by incoming analytes and lead to mass-transport effects, or introduce non-specific binding in certain applications. The HuProt Human Proteome Microarray is a multiplexed platform that requires robotic printing of all 21,000 proteins on its database from a pre-purified recombinant protein stock; however, expressing and purifying thousands of protein stocks, followed by printing individual 21,000 proteins on an array is not only cumbersome and time-consuming but causes loss in functionality in addition to being expensive and non-customizable. Importantly, HuProt arrays use end-point fluorescence detection and hence are incompatible with kinetic analysis.

To address the challenges of making functional protein microarrays in HTP with low-cost, quick turnaround, and which are readily customizable and allow for real-time, large-scale measurement of interaction kinetics, SPOC Proteomics, Inc. has developed a novel Sensor-Integrated Proteome On Chip technology (SPOC®) which automates the production and capture purification of *in situ* cell-free expressed functional protein arrays for quantitative kinetic analysis. To the best of our knowledge, SPOC is the only large-scale proteome array platform available today for large-scale study that overcomes the traditional challenges of protein microarray technologies highlighted here.

## Results and Discussion

### 1. SPOC as a novel platform for high throughput production of proteins on biosensor chips

SPOC is a first-of-its-kind proteomic biosensor system capable of delivering comprehensive quantitative, qualitative, and kinetic data within a single large-scale assay. We have successfully demonstrated *in situ* expression and simultaneous capture of proteins from immobilized plasmid DNAs directly onto a functionalized biosensor chip yielding a functional protein array (or SPOC array) for label-free analysis by SPR (**Fig. 1a**). The SPOC platform is scalable, versatile, and highly customizable. First, a customized plasmid DNA array is created by printing the plasmids onto a 25 x 75 mm silicon nanowell slide containing thousands of 2.0 nL volume nanowells, with center-to-center spacing of 225 to 375 µm for 30,000 wells/slide (30k) and 10,000 wells/slide (10k), respectively (**Fig. 1b**). The assay is currently configured such that proteins of interest are expressed from the printed plasmids as HaloTag fusion proteins, i.e. proteins fused with Halo protein as a common tag, for later covalent capture on the array. Next, the DNA-printed silicon nanowell slide is held in proximity to the surface of the biosensor capture slide whose surface has been pre-functionalized with HaloTag chloro-alkane linker to covalently capture the HaloTag protein, on the expressed fusion protein. A human HeLa-cell based, *in vitro* transcription and translation (IVTT) lysate mix is then injected between the nanowell slide and biosensor slides which are subsequently press-sealed against each other effectively isolating the nanowells to create nanoliter-volume reaction chambers for isolated protein expression (IPC). The sandwich is then incubated at 30°C for 2-4 hours during which the IVTT lysate expresses the protein encoded in the printed plasmid DNA contained in each nanowell, which are simultaneously capture-purified onto the biosensor capture slide forming the SPOC protein arrays. The entire process is facilitated by a custom AutoCap instrument developed and manufactured in house. This instrument contains four independent expression channels that enable four 25 mm x 75 mm nanowell slides to be expressed simultaneously, in a semi-automated fashion. From each nanowell slide, 4 SPR biosensors containing SPOC arrays can be produced, yielding 16 SPOC chips from four channels within a single expression run. Importantly, the number of proteins that can be arrayed on any capture slide is dependent on the size of the capture slide and density of the DNA-printed wells in the silicon nanowell slides. For example, our 10k silicon nanowell slide design expresses proteins in staggered pattern at a density of 716 protein spots/sq-cm which is equivalent to ∼1000 spots on the analysis area of the Carterra LSA^XT^ SPR instrument. However, our 30k nanowell slide design produces up to 1,975 spots/sq-cm, or 2400 protein spots on the Carterra LSA^XT^ SPR analysis area. **Figure 1c** shows an actual protein array on a gold biosensor produced by SPOC, highlighting distinct spots where individual proteins were immobilized. This innovative AutoCap instrument effectively reduces the cost by an astounding 10-100 times compared to traditional workflows that involve recombinant protein production and purification, making HTP proteomic screening more accessible.

**Figure 1.**
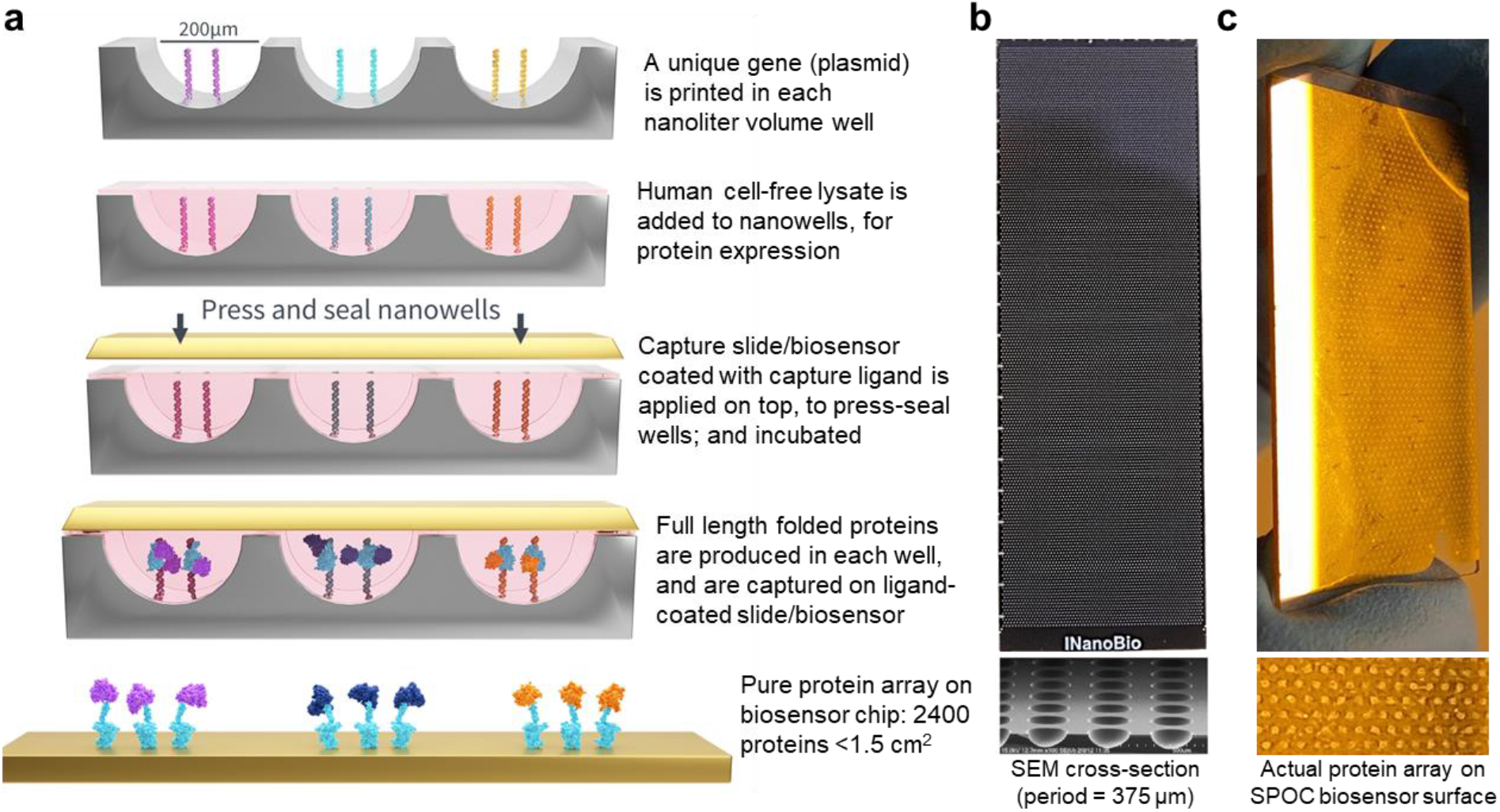
Schematic of *in situ* protein production in nanowell slide and capture on SPOC biosensor chip. **(a)** By leveraging human cell-free *in vitro* transcription and translation (IVTT) lysate, SPOC arrays are produced via *in situ* expression of full-length folded proteins, which are encoded by recombinant plasmid DNAs printed in nanowell slides. Expressed proteins are simultaneously capture-purified as a monolayer of arrayed spots onto biosensor chips for label-free analysis by SPR. The entire process from protein array production on biosensor chip to label-free kinetic analysis is called SPOC and is facilitated by our custom SPOC protein array production instrument. This instrument can produce 4 SPOC biosensors from a single nanowell slide. The instrument has four channels in each of which can produce a nanowell slide in a semi-automated fashion; ultimately yielding up to sixteen SPOC chips in a single run. (**b**) Image of a 10k nanowell slide into which expression plasmid DNAs are printed. Protein arrays are expressed from the nanowell slide are captured on a variety of substrates, including glass microscope slides for standard fluorescent screens, or gold-coated SPOC biosensors for label-free kinetic screening via SPR. A scanning electron microscope (SEM) image of a nanowell slide cross-section is shown below the full top-down image of the nanowell slide (25 mm x 75 mm). (**c**) Image of a SPOC biosensor and the protein microarray pattern formed on the surface of the gold-coated biosensor slide after the protein expression/capture process outlined in **a**.

### 2. Functional validation of high-density SPOC array using Carterra LSA^XT^ SPR instrument

SPOC technology is adaptable to any SPR instrument; however, for this publication, the Carterra LSA^XT^ SPR and Horiba Scientific OpenPleX SPR imaging (SPRi) instruments were used for validation because both instruments are amenable to HTP screening and require minimal effort to integrate with our SPOC protein array platform. Proteins captured on SPR biosensor chips for kinetic assay are henceforth referred to as protein-ligands. The functionality of the SPOC protein array is interrogated in this study using various screens including: detection of IVTT expressed proteins at discrete spots using protein-specific antibodies, differential binding kinetics of antibodies against variants of the same protein, demonstration of post translational modification (PTM) capabilities of arrayed proteins, autophosphorylation activity of arrayed proteins, and longitudinal stability of immobilized IVTT proteins after 10 days of slide storage.

We have demonstrated that SPOC arrays can be regenerated during SPR experiments, enabling multiple sample injections and kinetic screening from a single SPOC protein biosensor chip (**Supplemental Fig. S1**). The HaloTag-fusion proteins are covalently bound to the HaloTag chloro-alkane linker coated on the SPR biosensor chips, and we have found that harsh, 10 mM Glycine-HCl (pH 2.4) regeneration buffer conditions can be used for certain interactions without significant loss of biosensor integrity or interaction detection, while a 2 M NaCl buffer formulation is preferable and as effective for more sensitive binding interactions. SPOC is compatible with either condition depending on the analyte, and effectively strips non-covalently bound analytes from the covalently bound HaloTag-fusion proteins that constitute the SPOC array, enabling repeat sample injections for multiplexed applications on a single SPOC chip.

#### 2.1 Antibody Reactivity and HTP-Deep Kinetic Analysis of SPOC Array

Protein-specific antibodies were demonstrated to bind to their respective proteins on the SPOC array with high specificity. **Figure 2** shows the result of antibody assays performed on SPOC protein chips screened via the Carterra LSA^XT^, validating the ability to detect multiple proteins on the SPOC array in HTP while providing primary kinetic information. Locations of protein array spots on SPR biosensor chip are defined by region of interest (ROI) selection, when viewing the optical image of the SPOC protein chip docked on the SPR instrument. To enable mapping of protein array spots during ROI selection, a physical hockey-stick-shaped orientation marker was created on the nanowell slide that is imprinted on the SPR slide during expression/capture of the SPOC array and which is visually apparent on the SPR flowcell optical image (**Fig. 2a**). This is important because the nanowell pattern/spots on the sensor slide can be mapped in reference to this marker to identify the location of each protein on the SPOC array. The chip is probed with anti-HaloTag antibody that binds to Halo protein at all expression spots, to ensure successful immobilization and capture of the expressed HaloTag fused protein array. Due to software limitations, only 384 spots were initially selected and analyzed at one time. **Figure 2b** shows representative sensorgrams from 384 ROI/spots on the SPOC array, revealing successful capture of the HaloTag proteins expressed from the plasmids in the nanowell slide. Varying signal intensity (Rmax) between ligand containing spots reflects differential expression and capture levels between the SPOC array proteins. This is not unexpected as the parent plasmid DNA array encodes genes of proteins with different sizes and characteristics, and therefore, varied levels of expression. Different strategies are currently being explored to improve and normalize the protein expression and capture levels, including (1) printing higher concentration of DNA or using linearized plasmid DNA to increase expression levels and achieve self-normalization via capture surface saturation, (2) maximizing the capture efficiency of the biosensor chip by optimizing self-assembled monolayer and hydrogel surface conditions, and (3) encoding low-expressing genes on plasmids with high expression promoters optimized for IVTT expression. Data normalization techniques are also being explored whereby kinetic data obtained from reactive ligands are normalized to that of the ligand’s respective HaloTag Rmax signal. Low protein capture levels can limit sensitivity and analyte detection. However, protein capture density is preferable in many cases to reduce mass transport and avidity effects. Overall, similar kinetic rates and affinities can be estimated based on surfaces with varying ligand densities (Kamat and Rafique, 2017). Furthermore, within the current iteration of the SPOC array, bivalent analytes which can interact with two protein-ligands simultaneously (such as the antibodies utilized in this study) can produce avidity affects, when utilized as analytes in SPR assays which adds complexity to the measured kinetic data. It is acknowledged that the kinetics of the observed antibody-analyte interactions reported are confounded by the bivalency of the antibody analytes, and that the 1:1 binding modeling of the double-referenced kinetic traces provide an imprecise estimate of the interaction kinetics. While the kinetics reported here cannot be isolated from the analyte avidity, the kinetic dissociation constants are described as “affinity” for simplicity. Methods to eliminate avidity effects by reducing and controlling the density of the captured protein-ligands on the SPOC array such that only one of the two antibody paratopes can associate with a single protein-ligand, favoring a 1:1 interaction, form part of future SPOC technology development roadmap. This will be achieved by engineering and controlling the density of HaloTag chloro-alkane linker coated on SPR biosensor chips, thereby controlling the proteins on biosensor surface to avoid avidity effects.

**Figure 2.**
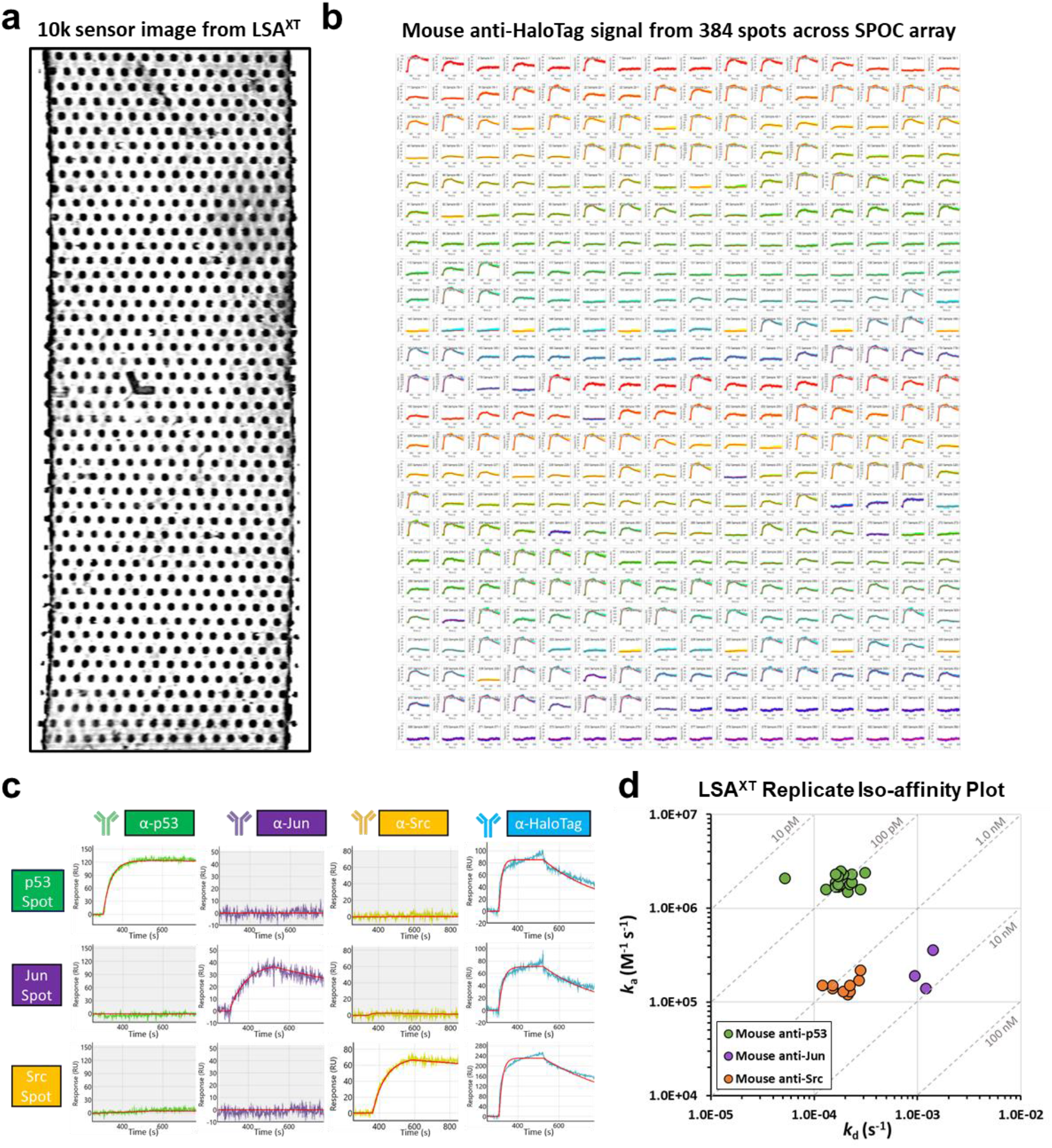
Carterra LSA^XT^ data of antibody interactions on SPOC chip. (**a**) Image of the SPOC sensor surface (10k array) as viewed from the flowcell of the LSA^XT^. The hockey-stick shaped orientation mark which aids in orienting and mapping the array is clearly visible in the center of the array. (**b**) Sensorgrams reporting the mouse anti-HaloTag antibody binding responses from across 384 spots simultaneously on the SPOC slide are shown and validate capture of IVTT expressed Halo fusion proteins. (**c**) Sensorgrams corresponding to each of three spots on the SPOC array containing p53, Jun, and Src HaloTag fusion proteins are shown demonstrating the binding responses observed after injection of mouse anti-p53 (16 nM), anti-Jun (67 nM), anti-Src (67 nM), and anti-HaloTag (133 nM) antibodies. The 1:1 interaction model is overlayed on the double-referenced data (red line). (**d**) Iso-affinity plot of the kinetics measured for the mouse anti-p53, anti-Jun, and anti-Src antibody injections (p53 = 20 spots across 3 sensors; Jun = 3 spots across 2 sensors; Src = 9 spots across 4 sensors).

Figure 2c emphasizes the highly specific detection of SPOC arrayed p53, Jun and Src when probed with their respective protein-specific antibodies. For example, when the sensor was probed with anti-p53 (first column), only the p53-HaloTag protein spot was detected. There was no response from spots where Jun and Src or other proteins were expressed and captured. Injection of anti-Jun (second column) showed response for the location of the sensor where Jun was captured, but not p53 and Src. Similarly, anti-Src could only detect Src proteins, highlighting the highly specific nature of SPOC assay. All spots showed response with anti-HaloTag. The surface of the sensor was regenerated with 10 mM Glycine-HCl (pH 2.4) between injections. Iso-affinity plot of the on-and off-rate measures estimated for the three commercial antibodies against their respective protein targets demonstrate the kinetic rates measured across multiple SPOC arrays are consistent (Fig. 2d). The iso-affinity plot clearly shows that the mouse anti-p53 antibody has a high affinity (avidity) with a KD = ∼100 pM while the mouse anti-Src spot replicates cluster around a KD = ∼1.0 nM. Compared to the anti-p53 antibody, the iso-affinity plot reveals that this lower affinity is driven by a slower on-rate, while the overall dissociation rate between the anti-p53 and anti-Src antibodies is comparable. The mouse anti-Jun antibody had the lowest measured affinity clustering around a *K*D just under 10 nM with a similar on-rate to that of the anti-Src antibody, but a comparably faster off-rate driving the lower affinity for the anti-Jun antibody. These results highlight the potential utility of the SPOC platform in antibody drug development leveraged for biological target validation (both on-and off-targets), and to screen artificial intelligence (AI) designed antibody clonal libraries to identify those with desired specificity and affinity.

#### 2.2 Ultra-High-Density SPOC arrays exhibit zero cross reactivity between adjacent protein spots

Figure 3a shows a SPOC protein chip image produced from our ultra-HTP 30k array which can yield up to 2400 protein spots on the Carterra LSA^XT^ analysis area with zero crosstalk between neighbor spots. The sole limitation of ultra-HTP SPOC analysis using the Carterra LSA^XT^ SPR instrument is the unavailability of software to display and analyze kinetic information from 2400 proteins at once. The current Carterra LSA^XT^ control and analysis software is capable of interrogating up to 384 spots, although a 1000 spot analysis software is in development and is expected to be commercially available in near future.

**Figure 3.**
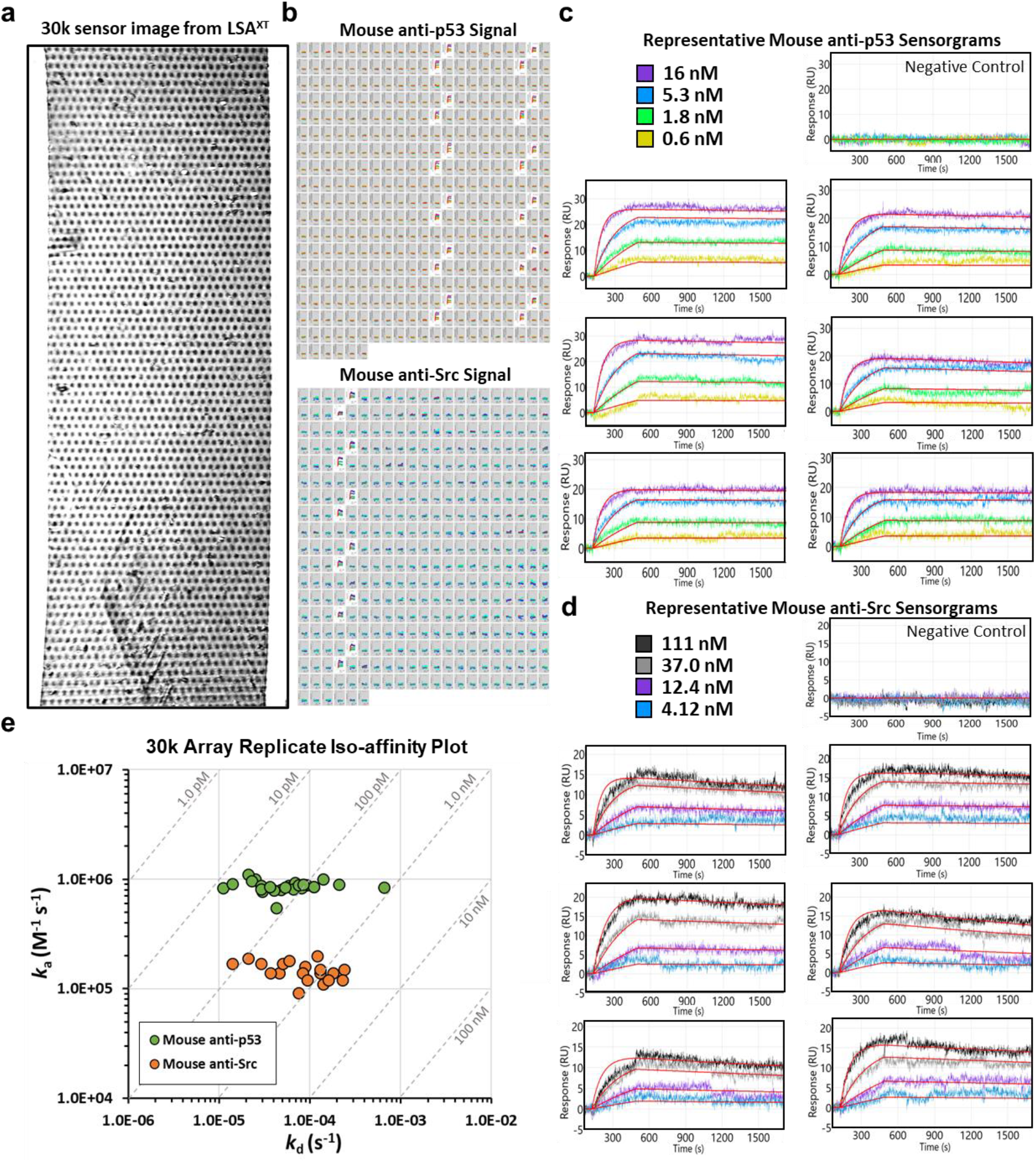
Carterra LSA^XT^ antibody kinetic data obtained from a SPOC chip produced from the high-density 30k nanowell slide. (**a**) Image of the SPOC sensor surface (30k array) as viewed from the flowcell of the LSA^XT^. (**b**) Sensorgrams from the Mouse anti-p53 and anti-Src antibody injections across 384 spots of NHv4 30k array are shown with positive binding responses unshaded, and areas with no binding responses shaded grey. Representative sensorgrams of the protein-specific antibodies binding to their specific protein targets (p53 or Src) are shown enlarged in **c** and **d**. (**c**) Six representative sensorgrams from a total of twenty-four p53-HaloTag spots bound by the p53 antibody from **b** are shown and the binding responses after the mouse anti-p53 antibody injections are displayed, the 1:1 interaction model (red lines) is overlayed on the double-referenced data. Concentrations of antibody titration are shown in the key above the sensorgrams alongside a sensorgram showing binding responses observed on a representative control spot of the array. (**d**) Six representative sensorgrams from a total of twelve Src-HaloTag spots are shown and the binding responses after the mouse anti-Src antibody injections are displayed, the 1:1 interaction model (red lines) is overlayed on the double-referenced data. (**e**) Iso-affinity plot of the kinetics measured for the mouse anti-p53 and anti-Src antibody injections (p53 = 26 spots across 2 sensors; Src = 19 spots across 2 sensors).

Zero cross reactivity (CR) between ultra-HTP 2400 protein spots was demonstrated by probing the 30k SPOC chip containing multiple p53 (24 spots) and Src (12 spots) protein spots with mouse anti-p53 and anti-Src antibodies (Fig. 3b). A larger version of the 384 ROI sensorgrams from the 30k array after anti-p53 and anti-Src antibody injection is reported in **Supplemental Figure S2** and **Supplemental Figure S3**, respectively. Reproducibility was ensured by injecting a serial titration of each detection antibody (four total injections each) followed by sensor regeneration with 10mM Glycine-HCl (pH = 2.37) between each antibody injection (Fig. 3c **& d**). Figure 3b shows highly specific detections of p53 and Src protein spots, indicated by an increase in their sensorgram response curves. Conversely, control spots around p53 and Src showed no response, indicating that protein expressed in sealed nanowells does not diffuse and capture at neighboring spots during the expression and capture step. This demonstrates close to zero crosstalk between spots. The iso-affinity plot of anti-p53 and anti-Src antibodies shows clustering of kinetic parameters from replicate spots, indicating minimal variations in on-rate estimates while the off-rate estimates were more variable (Fig. 3e). The variability in the off-rate estimates appear specific to the 30k array as compared to the kinetics measured on the 10k array (Fig. 2) and may be due to the challenges such small spots mount to precise ROI placement and could be improved with better reference ROI selection and placement during sensor installation.

#### 2.3 Analysis of SPOC Arrays using Orthogonal OpenPleX SPRi Instrument

Figure 4 shows the result of SPRi analysis using data obtained from a SPOC array on the lower throughput OpenPleX SPRi instrument (Horiba Scientific). The OpenPleX analysis area is approximately 1.0 sq-cm and therefore using our 10k silicon nanowell slide design we are able to interrogate up to 500 SPOC proteins (Fig. 4a). As expected, the orientation marker was visible on the flowcell image (Fig. 4a). In ‘SPR imaging’ based configurations like the OpenPleX instrument, the CCD camera which monitors the amount of reflected light from the sensor is fixed at an angle close to the SPR reflectance dip or minimum (the point at which most of the light is no longer reflected due to the surface-plasmon effect). Analyte binding to ligands on the sensor surface leads to a refractive index change that shifts the minimum and thus the camera records a concomitant increase in reflected light at the area of the sensor where binding is occurring, which is expressed as precent reflectivity in the sensorgrams. Due to this unique SPRi configuration, a difference image can be generated which shows the view from the CCD camera as analytes bind and produce a signal in the form of increased reflected light. At baseline, the difference image is black as it is set near the minimum and records little reflected light. In our case, as analytes bind to the ligand on the SPOC array, the spots themselves become visible in the difference image and interactions at discrete locations of the array can be visualized while the instrument records the magnitude of the percent reflectivity change occurring at each spot on the array. The signal visualized from the difference image can be directly correlated to the kinetic traces plotted in the sensorgrams generated from the OpenPleX. The SPOC array becomes plainly visible in the OpenPleX difference image after injection of 133 nM anti-HaloTag antibody which binds the HaloTag fusion proteins immobilized on the chip (Fig. 4b). The SPRi difference image showed positive detection of many HaloTagged proteins on the array. The varying intensity signals is consistent with our observation in Fig. 2b, highlighting differential expression and capture levels between SPOC proteins at expected locations based on the printed DNA (array map for the OpenPleX sensor is shown in **Supplemental Figure S4**). The OpenPleX SPRi instrument was also used to detect protein-specific antibody interaction on the SPOC array. Figures 4c and **4d** shows the difference image and sensorgrams, respectively, of two Jun spots on the SPOC array following interrogation with three concentrations of anti-Jun antibody (50, 100, and 200 nM). Figures 4e and **4f** depict similar information for four SPOC p53 protein spots after interrogation with 1.78 and 17.8 nM injections of anti-p53 antibody. These results show highly specific detection of SPOC proteins, with zero cross reactivity. The OpenPleX iso-affinity plot reveals minimal variation in computed kinetic parameters for the anti-Jun antibody, with higher variation for the anti-p53 kinetic measures between replicates (**Table 3**). Comparison of the KD values obtained from the manual injection-based OpenPleX instrument versus the highly automated and modern Carterra LSA^XT^ showed differences in measured kinetic parameters for the interactions and may be due to instrument features. The control/analysis software on our OpenPleX was dated and imaging-based SPR is generally less sensitive with a limit of detection ∼5-10 pg/mm^2^ compared to other configurations like angular-based instruments such as the Carterra LSA^XT^ which can be as low as 0.1 pg/mm^2^ (Homola et al., 1999; Wang et al., 2019). Discrepancies between the kinetic rates may also be due to the fact that the OpenPleX data were collected from bare gold sensors derivatized with Thiol-PEG-NHS self-assembled monolayers, whereas the SPOC arrays analyzed on the Carterra LSA^XT^ were performed on hydrogel HC30M slides (Xantec).

**Figure 4.**
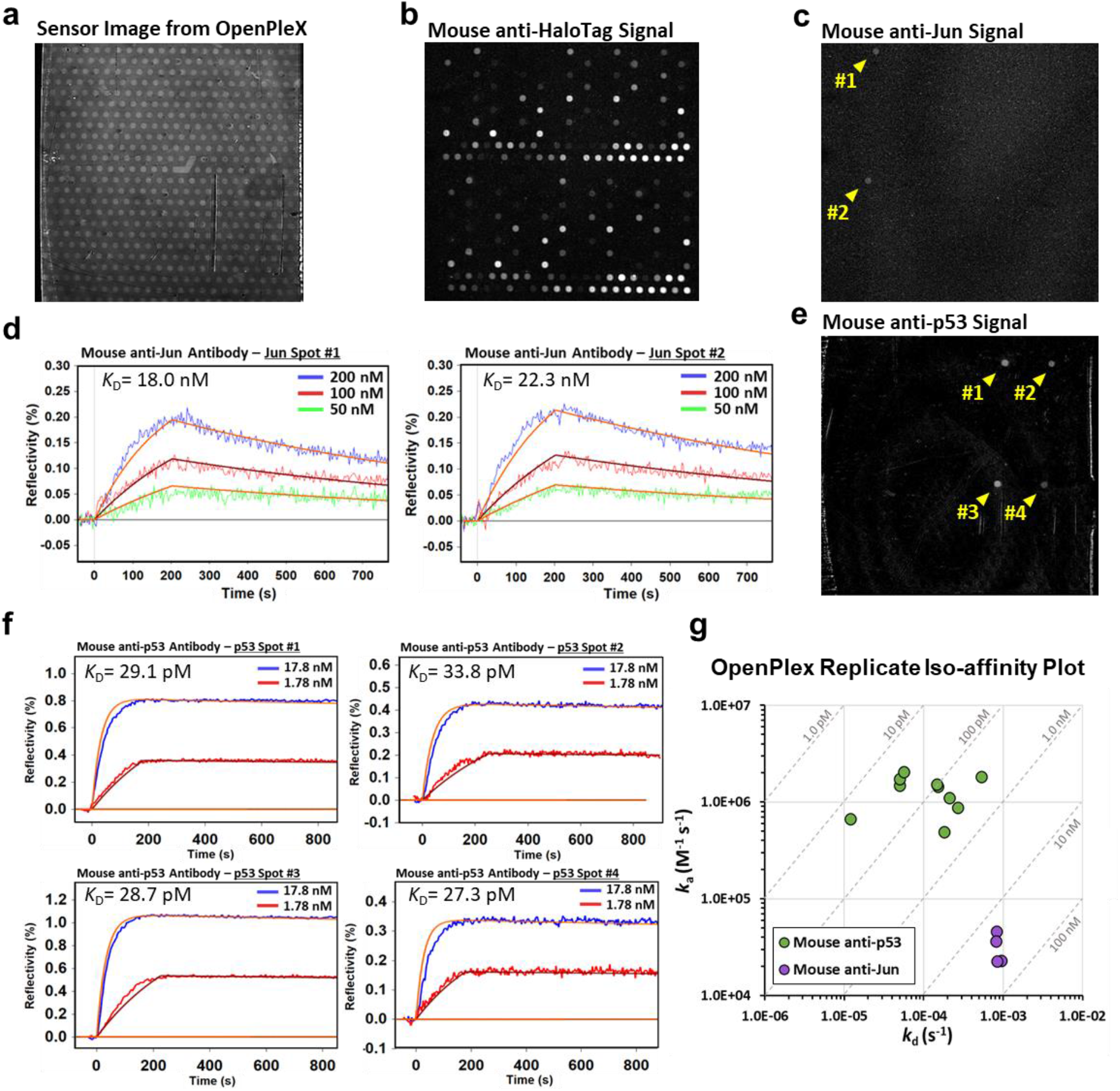
OpenPleX data of antibody interactions on SPOC chip. (**a**) Image of the SPOC sensor surface (10k pattern) as viewed from the flowcell of the OpenPleX. The hockey-stick shaped orientation mark is visible in the center of the array. (**b**) Difference image showing the pattern of binding observed on the SPOC biosensor surface after injection of mouse anti-HaloTag antibody (133 nM). (**c**) Difference image showing the pattern of binding observed after the Mouse anti-Jun antibody injection (200 nM) which specifically detects two spots (yellow marks) where Jun-HaloTag was capture-purified as expected. (**d**) 1:1 fit (orange line) OpenPleX data of the anti-Jun antibody binding to Jun Spots #1 and #2 from **c** show similar binding profiles and equilibrium disociation constants. (**e**) Difference image shows the pattern of binding observed after the mouse anti-p53 antibody injection (17.8 nM) which specifically detects four spots (yellow marks) where p53-HaloTag was capture-purified as expected. (**f**) 1:1 fit (orange lines) OpenPleX data of the anti-p53 antibody binding to the four p53 spots detected in **e** show similar binding profiles and equilibrium dissociation constants. (**g**) Iso-affinity plot of the kinetics measured for the Mouse anti-p53 and anti-Jun injections (p53 = 10 spots across 3 sensors; Jun = 4 spots across 2 sensors).

**Table 1.** 10k SPOC array kinetic parameters derived from three protein specific antibody analytes binding to their respective target ligands as measured on the Carterra LSA^XT^ instrument. Reported values were utilized to generate the iso-affinity plot in Fig. 2d.

| Analyte | SPOC Ligand | $k_a$ (M<sup>-1</sup>s<sup>-1</sup>) | $k_d$ (s<sup>-1</sup>) | $K_D$ (M) |
| --- | --- | --- | --- | --- |
| Mouse anti-p53 | p53 - Spot 1 | 1.8E+06 | 2.1E-04 | 1.17E-10 |
|  | p53 - Spot 2 | 1.7E+06 | 1.7E-04 | 1.00E-10 |
|  | p53 - Spot 3 | 1.6E+06 | 2.8E-04 | 1.75E-10 |
|  | p53 - Spot 4 | 1.8E+06 | 1.8E-04 | 1.00E-10 |
|  | p53 - Spot 5 | 1.8E+06 | 1.6E-04 | 8.89E-11 |
|  | p53 - Spot 6 | 1.9E+06 | 1.6E-04 | 8.42E-11 |
|  | p53 - Spot 7 | 1.6E+06 | 1.3E-04 | 8.13E-11 |
|  | p53 - Spot 8 | 1.5E+06 | 2.1E-04 | 1.40E-10 |
|  | p53 - Spot 9 | 1.7E+06 | 1.6E-04 | 9.41E-11 |
|  | p53 - Spot 10 | 1.8E+06 | 1.7E-04 | 9.44E-11 |
|  | p53 - Spot 11 | 1.8E+06 | 2.2E-04 | 1.22E-10 |
|  | p53 - Spot 12 | 1.9E+06 | 2.3E-04 | 1.21E-10 |
|  | p53 - Spot 13 | 2.4E+06 | 3.1E-04 | 1.29E-10 |
|  | p53 - Spot 14 | 2.2E+06 | 1.7E-04 | 7.73E-11 |
|  | p53 - Spot 15 | 2.1E+06 | 5.2E-05 | 2.48E-11 |
|  | p53 - Spot 16 | 2.3E+06 | 2.3E-04 | 1.00E-10 |
|  | p53 - Spot 17 | 2.3E+06 | 1.9E-04 | 8.26E-11 |
|  | p53 - Spot 18 | 2.5E+06 | 1.8E-04 | 7.20E-11 |
|  | p53 - Spot 19 | 2.2E+06 | 1.7E-04 | 7.73E-11 |
|  | p53 - Spot 20 | 2.3E+06 | 1.6E-04 | 6.96E-11 |
| Mouse anti-Jun | Jun - Spot 1 | 3.6E+05 | 1.4E-03 | 3.89E-09 |
|  | Jun - Spot 2 | 1.4E+05 | 1.2E-03 | 8.57E-09 |
|  | Jun - Spot 3 | 1.9E+05 | 9.4E-04 | 4.95E-09 |
| Mouse anti-Src | Src - Spot 1 | 1.7E+05 | 2.7E-04 | 1.59E-09 |
|  | Src - Spot 2 | 2.2E+05 | 2.8E-04 | 1.27E-09 |
|  | Src - Spot 3 | 1.2E+05 | 2.1E-04 | 1.75E-09 |
|  | Src - Spot 4 | 1.4E+05 | 1.5E-04 | 1.07E-09 |
|  | Src - Spot 5 | 1.3E+05 | 2.2E-04 | 1.69E-09 |
|  | Src - Spot 6 | 1.3E+05 | 1.9E-04 | 1.46E-09 |
|  | Src - Spot 7 | 1.5E+05 | 1.5E-04 | 1.00E-09 |
|  | Src - Spot 8 | 1.5E+05 | 2.2E-04 | 1.47E-09 |
|  | Src - Spot 9 | 1.5E+05 | 1.2E-04 | 8.00E-10 |

**Table 2.** 30k SPOC array kinetic parameters derived from two protein specific antibody analytes binding to their respective target ligands as measured on the Carterra LSA^XT^ instrument. Reported values were utilized to generate the iso-affinity plot in Fig. 3e.

| Analyte | SPOC Ligand | $k_a$ (M<sup>-1</sup>s<sup>-1</sup>) | $k_d$ (s<sup>-1</sup>) | $K_D$ (M) |
| --- | --- | --- | --- | --- |
| Mouse anti-p53 | p53 - Spot 1 | 8.4E+05 | 6.6E-04 | 7.86E-10 |
|  | p53 - Spot 2 | 1.0E+06 | 2.5E-05 | 2.50E-11 |
|  | p53 - Spot 3 | 7.7E+05 | 3.0E-05 | 3.90E-11 |
|  | p53 - Spot 4 | 8.7E+05 | 5.8E-05 | 6.67E-11 |
|  | p53 - Spot 5 | 5.5E+05 | 4.3E-05 | 7.82E-11 |
|  | p53 - Spot 6 | 8.5E+05 | 6.3E-05 | 7.41E-11 |
|  | p53 - Spot 7 | 8.5E+05 | 1.1E-04 | 1.29E-10 |
|  | p53 - Spot 8 | 8.3E+05 | 1.1E-05 | 1.33E-11 |
|  | p53 - Spot 9 | 8.0E+05 | 4.8E-05 | 6.00E-11 |
|  | p53 - Spot 10 | 9.3E+05 | 6.9E-05 | 7.42E-11 |
|  | p53 - Spot 11 | 8.8E+05 | 9.2E-05 | 1.05E-10 |
|  | p53 - Spot 12 | 1.1E+06 | 2.1E-05 | 1.91E-11 |
|  | p53 - Spot 13 | 9.6E+05 | 2.3E-05 | 2.40E-11 |
|  | p53 - Spot 14 | 7.9E+05 | 4.0E-05 | 5.06E-11 |
|  | p53 - Spot 15 | 8.3E+05 | 8.1E-05 | 9.76E-11 |
|  | p53 - Spot 16 | 8.5E+05 | 1.1E-04 | 1.29E-10 |
|  | p53 - Spot 17 | 8.7E+05 | 2.9E-05 | 3.33E-11 |
|  | p53 - Spot 18 | 8.1E+05 | 6.6E-05 | 8.15E-11 |
|  | p53 - Spot 19 | 1.0E+06 | 1.4E-04 | 1.40E-10 |
|  | p53 - Spot 20 | 8.8E+05 | 7.5E-05 | 8.52E-11 |
|  | p53 - Spot 21 | 8.9E+05 | 8.5E-05 | 9.55E-11 |
|  | p53 - Spot 22 | 9.1E+05 | 1.4E-05 | 1.54E-11 |
|  | p53 - Spot 23 | 9.0E+05 | 2.1E-04 | 2.33E-10 |
|  | p53 - Spot 24 | 8.4E+05 | 5.3E-05 | 6.31E-11 |
|  | p53 - Spot 25 | 8.1E+05 | 2.9E-05 | 3.58E-11 |
|  | p53 - Spot 26 | 8.5E+05 | 3.6E-05 | 4.24E-11 |
| Mouse anti-Src | Srs - Spot 1 | 1.4E+05 | 1.8E-04 | 1.29E-09 |
|  | Srs - Spot 2 | 1.4E+05 | 1.3E-04 | 9.29E-10 |
|  | Srs - Spot 3 | 1.6E+05 | 8.9E-05 | 5.56E-10 |
|  | Srs - Spot 4 | 1.7E+05 | 1.4E-05 | 8.24E-11 |
|  | Srs - Spot 5 | 1.4E+05 | 8.3E-05 | 5.93E-10 |
|  | Srs - Spot 6 | 1.7E+05 | 2.9E-05 | 1.71E-10 |
|  | Srs - Spot 7 | 1.4E+05 | 4.6E-05 | 3.29E-10 |
|  | Srs - Spot 8 | 1.9E+05 | 2.1E-05 | 1.11E-10 |
|  | Srs - Spot 9 | 1.5E+05 | 1.3E-04 | 8.67E-10 |
|  | Srs - Spot 10 | 1.4E+05 | 3.7E-05 | 2.64E-10 |
|  | Srs - Spot 11 | 1.7E+05 | 5.0E-05 | 2.94E-10 |
|  | Srs - Spot 12 | 9.1E+04 | 7.5E-05 | 8.24E-10 |
|  | Srs - Spot 13 | 1.5E+05 | 2.4E-04 | 1.60E-09 |
|  | Srs - Spot 14 | 1.2E+05 | 2.3E-04 | 1.92E-09 |
|  | Srs - Spot 15 | 2.0E+05 | 1.2E-04 | 6.00E-10 |
|  | Srs - Spot 16 | 1.1E+05 | 1.4E-04 | 1.27E-09 |
|  | Srs - Spot 17 | 1.2E+05 | 1.6E-04 | 1.33E-09 |
|  | Srs - Spot 18 | 1.8E+05 | 5.9E-05 | 3.28E-10 |
|  | Srs - Spot 19 | 1.2E+05 | 9.4E-05 | 7.83E-10 |

**Table 3.** 10k SPOC array kinetic parameters derived from two protein specific antibody analytes binding to their respective target ligands as measured on the OpenPleX instrument. Reported values were utilized to generate the iso-affinity plot in Fig. 4g.

| Analyte | SPOC Ligand | $k_a$ ($M^{-1}s^{-1}$) | $k_d$ ($s^{-1}$) | $K_D$ (M) |
| --- | --- | --- | --- | --- |
| Mouse anti-p53 | p53 - Spot 1 | 1.8E+06 | 5.3E-05 | 2.91E-11 |
|  | p53 - Spot 2 | 1.5E+06 | 5.0E-05 | 3.38E-11 |
|  | p53 - Spot 3 | 1.7E+06 | 5.0E-05 | 2.87E-11 |
|  | p53 - Spot 4 | 2.1E+06 | 5.6E-05 | 2.73E-11 |
|  | p53 - Spot 5 | 1.1E+06 | 2.1E-04 | 1.91E-10 |
|  | p53 - Spot 6 | 4.9E+05 | 1.8E-04 | 3.67E-10 |
|  | p53 - Spot 7 | 8.8E+05 | 2.7E-04 | 3.07E-10 |
|  | p53 - Spot 8 | 6.7E+05 | 1.2E-05 | 1.79E-11 |
|  | p53 - Spot 9 | 1.4E+06 | 1.5E-04 | 1.04E-10 |
|  | p53 - Spot 10 | 1.5E+06 | 1.5E-04 | 9.61E-11 |
| Mouse anti-Jun | Jun - Spot 1 | 4.59E+04 | 8.30E-04 | 1.81E-08 |
|  | Jun - Spot 2 | 3.64E+04 | 8.10E-04 | 2.23E-08 |
|  | Jun - Spot 3 | 2.30E+04 | 9.50E-04 | 4.13E-08 |
|  | Jun - Spot 4 | 2.27E+04 | 8.40E-04 | 3.70E-08 |

Detection of post-translational citrullination of four HIST1H3A protein spots on the SPOC array was demonstrated with OpenPleX instrument. Prior to on-slide citrullination, lack of citrulline group on the HIST1H3A protein spots was confirmed by probing a freshly prepared SPOC array with anti-cit-HIST1H3A, which also contained few protein spots not known to be citrullinated. As expected, after the citrullination reaction and injection of anti-cit-HIST1H3A and secondary antibody, the resulting SPRi difference image and graph of reflectivity change showed marked increase in HIST1H3A spots contrast and reflectivity signals (**Supplemental Fig. S5**; green boxes). Conversely, the control protein spots showed no signals, underscoring the potential of the SPOC platform for use in studying certain PTMs. This assay is currently being optimized to output primary kinetic information.

### 3. Orthogonal immunofluorescence validation of SPOC protein arrays captured on glass slides

The production of SPOC arrays is not limited to biosensor chips alone, but also possible on a standard 25 x 75 mm microscope glass slide to yield high-density protein arrays of up to 30,000 proteins for semi-quantitative analysis by immunofluorescence (Fig. 5a**-c**). The functionality of the SPOC protein array on glass was validated via series of experiments using an array containing replicates spots of HaloTagged p53, Fos, Jun, FGA, HIST1H3A, among many others. Protein expression/capture were confirmed by HaloTag antibody immunofluorescence assay (Fig. 6a**&b**). The SPOC array exhibited minimal cross reactivity (between adjacent protein spots (Fig. 6c). Cross reactivity (CR) was determined by dividing the average anti-p53 antibody signal in wells surrounding each of the 184 total p53 spots with the average anti-p53 signal in p53 spots. The variation (CV) was also measured by dividing the standard deviation of the fluorescent intensity of all p53 spots by the average fluorescent intensity of the same spots and multiplying by 100 and was calculated to be 31% for the anti-Halo-probed slide shown in Fig. 6c, and 28% for all four dense areas shown enlarged on Fig. 6d, yet, spots are still very clearly observable via antibody stain.

**Figure 5.**
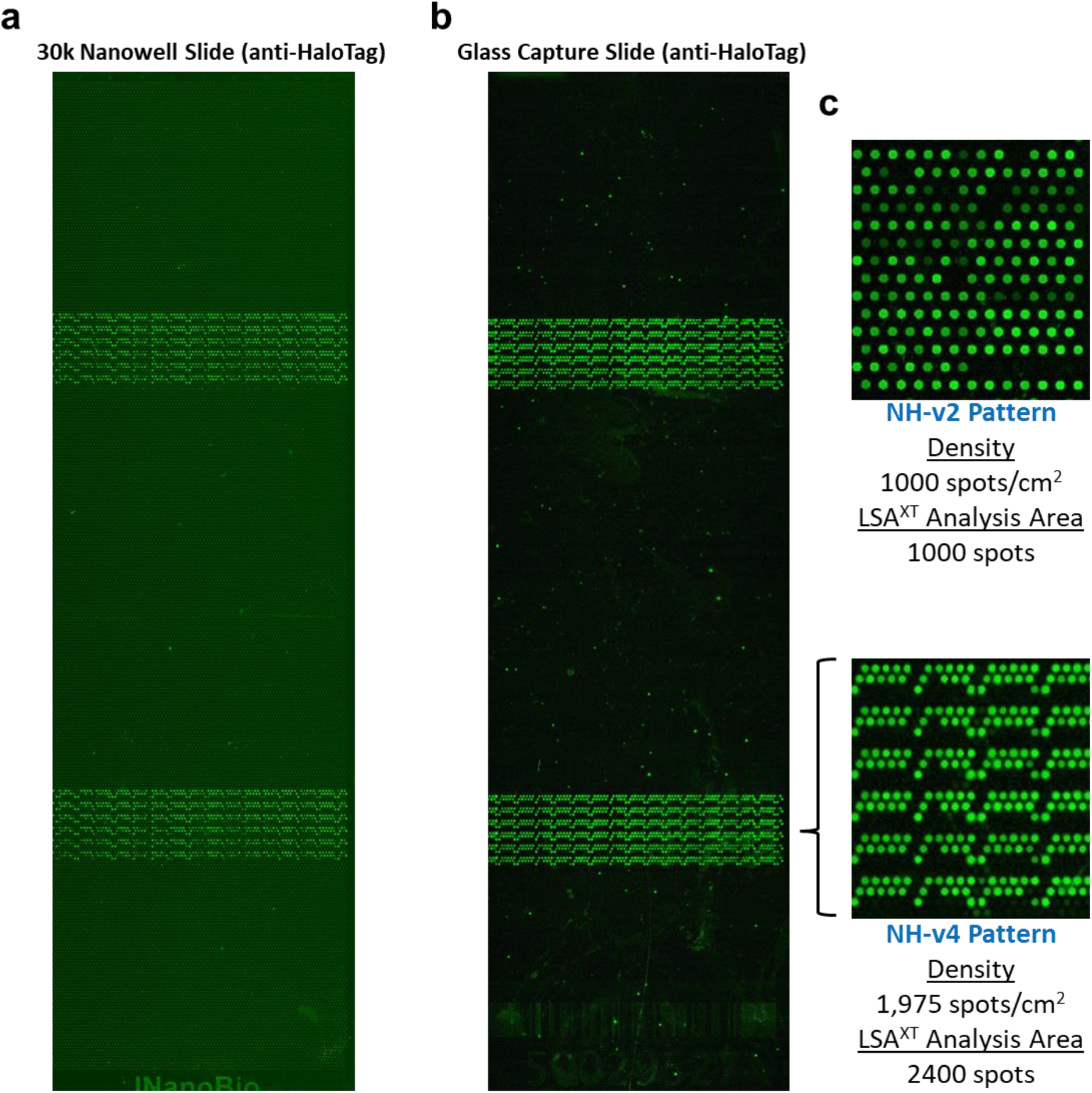
Nanowell slide design facilitates high-density SPOC arrays capture-purification. (**a**) Mouse anti-HaloTag fluorescence staining pattern observed on a high-density 30k nanowell slide design demonstrates expression of HaloTag fusion proteins in the nanowells of two subarrays containing various expression plasmids of interest. (**b**) Mouse anti-HaloTag fluorescence staining pattern observed on a glass capture slide functionalized with HaloTag chloro-alkane linker reveals diffusion-free capture of the high-density protein array generated from the 30k nanowell slide after capture-purification on the SPOC instrument (**c**) A comparison of the spot patterning achieved on the 10k versus the 30k nanowell slides, and the respective spot densities achieved is shown. The 30k design generates a >2-fold denser array than the 10k.

**Figure 6.**
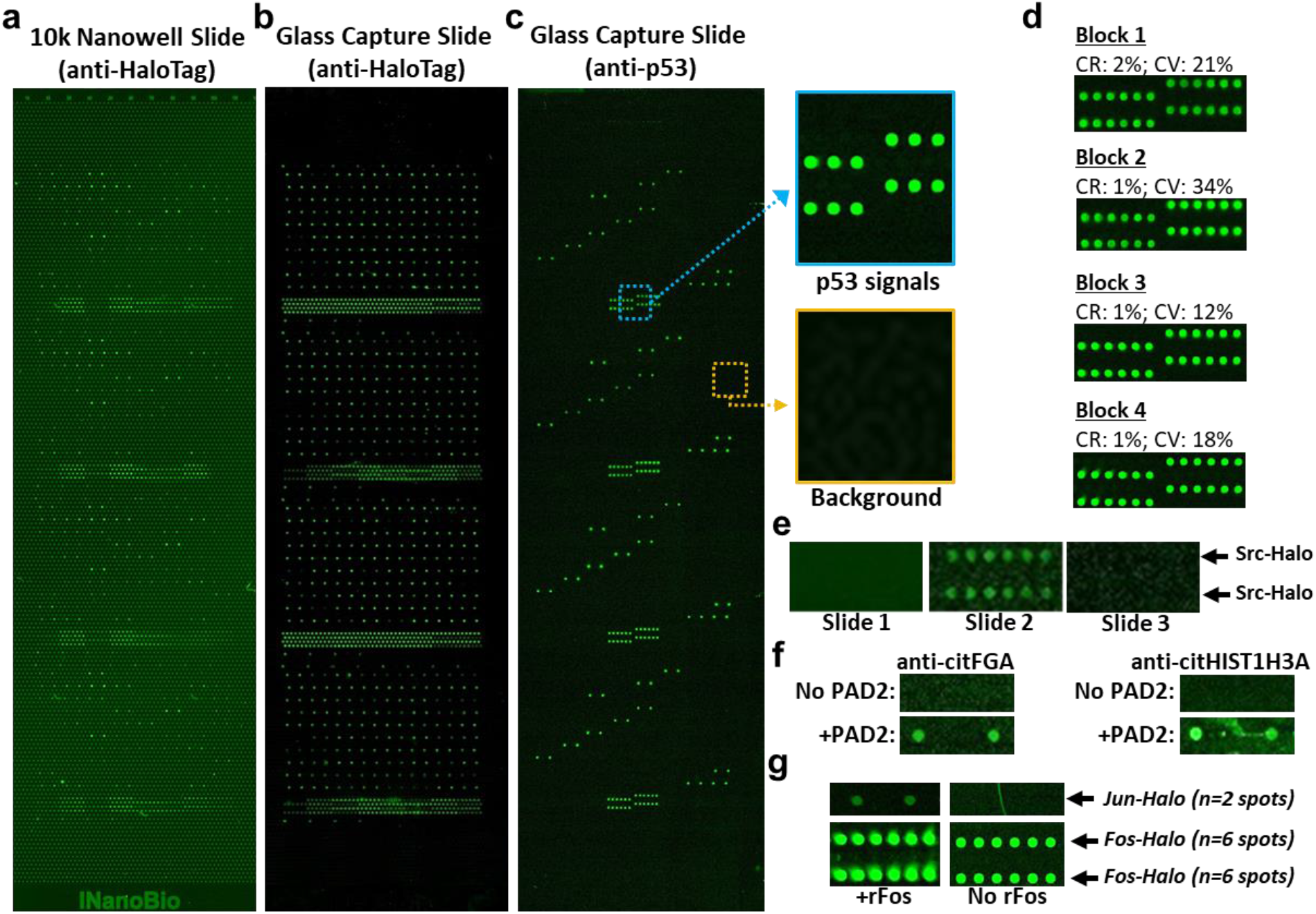
Automated production of pure in-situ protein arrays capture-purified on glass. (**a**) Validation data showing fluorescence images of Halo-tagged proteins expressed in silicon nanowell, probed with anti-Halo, (**b**) captured onto HaloTag chloro-alkane linker-modified capture glass slides which were probed with anti-Halo, and (**c**) the same slide layout as shown in (**b**), but probed with anti-p53 antibody to detect p53 protein spots. These images show strong HaloTag and p53 signals on capture-purified protein arrays on capture glass slide, with <2% crosstalk between proteins in neighboring spots where protein was otherwise detected in (**b**). (**d**) Shows statistical analysis of cross reactivity (CR) and variation (CV) of p53 spot signals calculated from each block as a whole (sparse and dense areas). The enlarged images show the dense area only of each block to demonstrate lack of signal between spots and in neighboring non-p53 spots. Functional validation testing of protein arrays captured on glass slides. (**e**) demonstrates the enzymatic capability of bound Src kinase proteins via on-slide phosphorylation and dephosphorylation. Slide 1-3 were stained with anti-phosphotyrosine (n = 12 Src spots). Slide 1 consisting freshly expressed and captured Halo-SRC kinase protein spots was dephosphorylated by incubating with alkaline phosphatase and then stained. Slide 2 - Src spots were phosphorylated by incubating with kinase buffer to allow Src to autophosphorylate, followed by anti-phosphotyrosine staining. Slide 3 – Src was dephosphorylated again followed by staining. (**f**) Proteins can be altered with known PTMs associated with autoimmunity, such as citrullination. Here, Halo-fibrinogen (FGA) and Halo-Hist1H3A spots were treated with or without PAD2 enzyme to catalyze citrullination and then probed with citrullination-specific anti-cit-FGA and anti-cit-HIST1H3A. (**g**) Capture protein can bind to known protein interaction partners, such as Fos and Jun. Halo-Fos and Halo-Jun spots were incubated with (**g**) or without (**h**) recombinant Fos in solution. Slides were then probed with anti-Fos to determine if Fos bound to Halo-Jun. Fos was detected on Halo-Jun spots only if first incubated with rFos, and was detected at all spots known to contain Halo-Fos as a positive control – demonstration functionality of produced capture-purified protein arrays.

In terms of functionality, the SPOC protein array on glass was shown to bind protein specific antibodies after probing with anti-p53, anti-Fos, anti-cit-FGA, and anti-cit-HIST1H3A antibodies. These antibodies specifically bound to only their protein targets on the SPOC array and were detected with a secondary fluorescently-tagged antibody. Fig. 6c shows a representative image of p53 fluorescence signals after probing with anti-p53, emphasizing nearly zero background CR. Further, the catalytic activity function of enzymes displayed on SPOC array was demonstrated using Src kinase as a model. This protein is capable of self (auto) phosphorylation on several of its tyrosine residues. To show activity, the SPOC array was first dephosphorylated and probed with anti-phosphotyrosine antibody (Fig. 6e, Slide 1). No signal was observed, indicating all phosphate groups were removed. On a matching slide, the SPOC array was dephosphorylated and then incubated in kinase buffer containing ATP. Phosphorylation of Src protein on this SPOC glass slide (Slide 2) was confirmed after staining with anti-phosphotyrosine antibody, as expected (Fig. 6.). By contrast, if the SPOC glass slide was treated once more with dephosphorylation buffer after kinase buffer incubation, no signal was observed from the anti-phosphotyrosine antibody (Fig. 6e, Slide 3). Further, post-translational modification of SPOC proteins was demonstrated by *in vitro* citrullination of HIST1H3A and FGA proteins on a SPOC glass slide array using PAD2 enzyme. The results show distinct anti-cit-FGA and anti-cit-HIST1H3A in test slides which were absent in the control slides incubated in the same conditions minus PAD2 (Fig. 6f). Finally, protein-protein interaction functionality of SPOC proteins was demonstrated using Jun proteins arrayed on a SPOC glass slide array (Fig. 6g). The slide was probed with or without a recombinant Fos protein in solution and then detected with anti-Fos antibody. The result revealed highly specific Fos signals on the two Jun protein spots (Fig. 6g left panel) as well as Fos protein spots (positive control), but no Fos signal was detected on the Jun protein spots in the control slide that was not probed with recombinant Fos (Fig. 6g right panel). Furthermore, the above Fos-Jun interactions, capacity to introduce citrulline PTM to HIST1H3A, in addition to monitoring of antibody-antigen interactions were observed to be stable and readily performed successfully after the SPOC arrays had been stored for 10 days under optimized conditions (**Supplemental Figure S6**). These results underscore the SPOC platform as a potential tool to study protein interactions, PTM, and protein activity assays in HTP.

### 5. SPOC platform for drug screening: Kinetic profiling of differential antibody response to SARS-CoV-2 Spike RBD variants

Considering the rapidly mutating SARS-CoV-2 virus responsible for the COVID-19 pandemic and the need to develop platform technologies for assessing the potency of neutralizing antibody drugs for variants of concern (VoC), we sought to validate if SPOC can resolve differential kinetics of antibody drug binding to variants of SARS-CoV-2 receptor binding domain (RBD) using the Carterra LSA^XT^ for SPR analysis. In addition to the wildtype Wuhan RBD, nine variants - Alpha, Delta, and Omicron variants and subvariants BA.1, BA.5, BA.2.75, BQ.1, XBB.1.16, XBB.1.5 and BQ.1.1 were arrayed on a SPOC chip and then screened for strength of binding to a commercial anti-RBD antibody (Proteintech Group, cat#: 67758-1) from two different manufacturer lots. Results of orthogonal qualitative anti-HaloTag fluorescence manual spotting assay revealed comparable protein expression levels profiles among the RBD variants as well as qualitative differential binding of the anti-RBD antibody. This antibody bound robustly to Wuhan, Alpha, and Delta RBD variants, but exhibited significantly lower binding affinity to Omicron BA1 (Fig. 7a) and binding was undetectable with other variants (data not shown).

**Figure 7.**
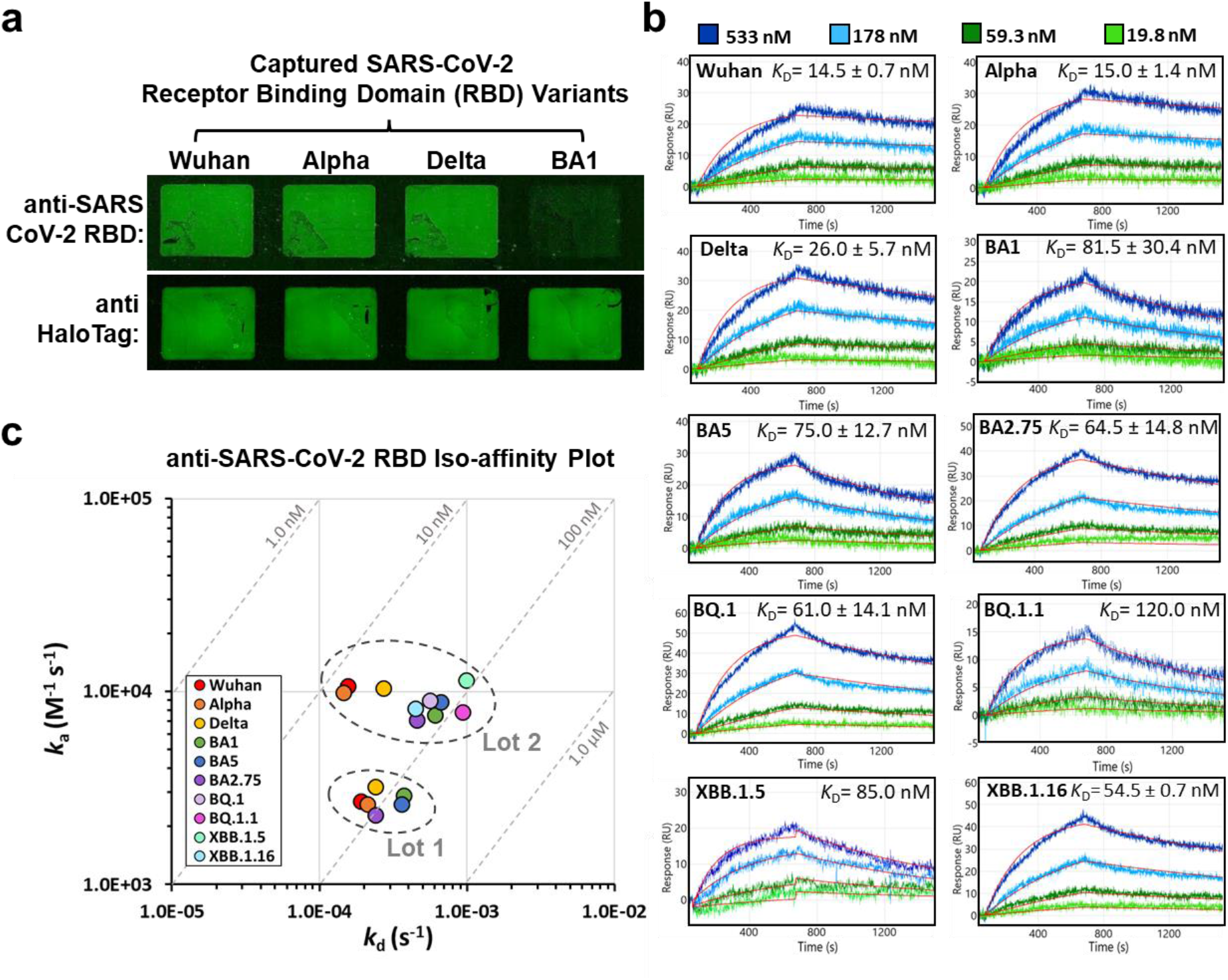
Carterra LSA^XT^ screening of mouse anti-SARS-CoV-2 receptor binding domain (RBD) against RBD variants arrayed on SPOC chip. (**a**) Orthogonal fluorescent glass-slide assay shows the staining pattern observed after capturing via manually spotting the indicated IVTT expressed SARS-CoV-2 RBD-HaloTag proteins and probing with mouse anti-RBD and HaloTag antibodies. (**b**) Sensorgrams reporting the binding responses observed after a serial titration of mouse anti-RBD antibody (Lot 2) was injected over a SPOC array containing 10 RBD VoC via the Carterra LSA^XT^. 1:1 fits are overlayed on the data (red curves) and the equilibrium dissociation constants were determined from duplicate screens from Lot 2 of the antibody. (**c**) Iso-affinity plot of the various kinetics measured for the mouse anti-RBD antibody from two different lots against the RBD VoC are reported. The antibody from Lot 1 was only screened on a sensor with 6 of the RBD VoC while Lot 2 was screened across 2 sensors containing all 10 CoV-2 RBDs of interest.

As expected, label-free screening by SPOC SPR provided additional information that was lacking in the end-point fluorescence assay, with real-time SPR capable of detecting binding of the anti-RBD antibody to all the RBD variants revealing differential kinetics, including variants that were undetected using fluorescence assay (e.g. BA1). In the first SPOC SPR assay, a SPOC chip containing five RBD variants and wildtype Wuhan was probed with Proteintech anti-RBD from manufacturer Lot 1. Evaluation of *K*D values revealed that Wuhan, Delta, and Alpha RBDs exhibited the strongest affinities to this antibody (*K*D = 70, 74, and 80 nM, respectively). These affinities were ∼2-fold higher than *K*D of Omicron variants BA.1 (127 nM), BA.5 (137 nM) and BA.2.75 (105 nM) and characterized by faster off-rates (**Supplemental Fig. S7** and Fig. 7c). The primary anti-RBD was highly specific, and did not bind to hundreds of other Halo-fusion proteins captured on the SPOC array (not shown).

In another set of experiments, a SPOC array of all nine SARS-CoV-2 RBD variants and Wuhan wildtype was probed with the Proteintech anti-RBD from manufacturer Lot 2. KD values of the Wuhan, Alpha and Delta RBD variants were KD of 15, 14, and 22 nM, respectively. The Omicron variants BA.1, BA.5, and BA.2.75 clustered together with values of 60, 66, and 75 nM, respectively. The KD values of other variants - BQ.1, XBB.1.16, XBB.1.5 were calculated as 71, 54, and 85 nM, respectively. The response curve of anti-RBD to BQ.1.1 variant was too low in this experiment and not computed by the analysis software. A repeat experiment was performed with the Lot 2 anti-RBD antibody using a replicate SPOC chip produced and screened one week apart. The result remained consistent, with the antibody exhibiting the highest affinity to Wuhan, Alpha and Delta RBD (mean KD ± standard deviation from both experiment with Lot 2; 14.5 ± 0.7, 15 ± 1.4, and 26 ± 5.7 nM, respectively). The mean KD values for BA.1, BA.5, BA.2.75, BQ.1, and XBB.1.16 variants, were found to be 81.5 ± 30.4, 75 ± 12.7, 64.5 ± 14.8, 61 ± 14.1, and 54.5 ± 0.7, respectively (Fig. 7b). In this repeat experiment, binding of the antibody against the XBB.1.5 variant was too low to extract kinetic information. The lowest anti-RBD binding affinity was obtained against the BQ.1.1 variant (KD = 120 nM). The minimal SD observed in these replicate SPOC screening of the Lot 2 anti-RBD validates the reproducibility of the SPOC assay. An iso-affinity plot of the estimated anti-RBD kinetics against the RBD VoC from the two antibody lots is depicted in Fig. 7c. The result shows the trend of differential binding kinetics similar in both lots, albeit with variations in KD values (Fig. 7c). In both screening assays, Wuhan, Alpha, and Delta RBD, which had the highest affinities, clustered together, while the other variants induced lower affinity binding (**Table 4**).

**Table 4.** Kinetic parameters derived from the Proteintech mouse anti-SARS-CoV-2 RBD antibody analyte binding to Wuhan RBD and variants. Reported values were utilized to generate the iso-affinity plot in Fig. 7c. Cells with dashes indicate interactions which were not measured, while cells with N/A indicate interactions where no kinetic parameters could be estimated due to low binding signal. The assay was performed on Carterra LSA^XT^.

| anti-RBD Ab Lot | Replicate | Kinetic Measure | Wuhan | Alpha | Delta | BA1 | BA5 | BA2.75 | BQ.1 | BQ.1.1 | XBB.1.5 | XBB.1.16 |
| --- | --- | --- | --- | --- | --- | --- | --- | --- | --- | --- | --- | --- |
| Lot 1 | #1 | $k_a$ (M<sup>-1</sup>s<sup>-1</sup>) | 2.7E+03 | 2.60E+03 | 3.20E+03 | 2.90E+03 | 2.60E+03 | 2.30E+03 | --- | --- | --- | --- |
| | | $k_d$ (s<sup>-1</sup>) | 1.9E-04 | 2.10E-04 | 2.40E-04 | 3.70E-04 | 3.60E-04 | 2.40E-04 | --- | --- | --- | --- |
| | | $K_D$ (M) | 7.0E-08 | 8.1E-08 | 7.5E-08 | 1.3E-07 | 1.4E-07 | 1.0E-07 | --- | --- | --- | --- |
| Lot 2 | #1 | $k_a$ (M<sup>-1</sup>s<sup>-1</sup>) | 1.3E+04 | 1.20E+04 | 1.20E+04 | 8.00E+03 | 9.20E+03 | 7.00E+03 | 9.80E+03 | N/A | 1.20E+04 | 8.70E+03 |
| | | $k_d$ (s<sup>-1</sup>) | 2.0E-04 | 1.70E-04 | 2.70E-04 | 4.80E-04 | 6.10E-04 | 5.20E-04 | 7.00E-04 | N/A | 9.80E-04 | 4.70E-04 |
| | | $K_D$ (M) | 1.5E-08 | 1.4E-08 | 2.3E-08 | 6.0E-08 | 6.6E-08 | 7.4E-08 | 7.1E-08 | N/A | 8.2E-08 | 5.4E-08 |
| | #2 | $k_a$ (M<sup>-1</sup>s<sup>-1</sup>) | 8.4E+03 | 7.70E+03 | 8.90E+03 | 7.10E+03 | 8.50E+03 | 7.20E+03 | 8.20E+03 | 7.80E+03 | N/A | 7.60E+03 |
| | | $k_d$ (s<sup>-1</sup>) | 1.1E-04 | 1.20E-04 | 2.70E-04 | 7.30E-04 | 7.20E-04 | 3.90E-04 | 4.20E-04 | 9.30E-04 | N/A | 4.20E-04 |
| | | $K_D$ (M) | 1.3E-08 | 1.6E-08 | 3.0E-08 | 1.0E-07 | 8.5E-08 | 5.4E-08 | 5.1E-08 | 1.2E-07 | N/A | 5.5E-08 |

Overall, the Proteintech anti-RBD antibody from Lot 2 demonstrated notably greater affinity compared to Lot 1, underscoring the SPOC platform’s potential for evaluating batch-to-batch variation in GMP-manufactured antibody products. Although the exact epitope of the Proteintech anti-RBD used in this experiment is unknown, the fact that the mutations present in the Omicron variant RBD significantly reduces antibody recognition may suggest that the epitope is closer to the N-terminus of the domain where the mutations on Omicron least overlap with those present in the other variants. Therefore, these findings also highlight the potential value of the SPOC platform in pandemic response situations, particularly when there is a need to promptly monitor and characterize the impact of mutations at immunogenic viral epitopes, or evaluate the potential risk of immune escape or diminished vaccine efficacy associated with any emerging novel variant.

### Conclusion

SPOC is a first-of-kind platform for high throughput proteomic kinetic profiling on biosensors, providing real-time label-free quantitative, qualitative, and kinetic affinity data. SPOC protein chips are rapidly customizable, contain folded full-length proteins, and enables 10-100 times reduced cost of assay due to the lack of requirement for laborious and expensive individual protein expression, pre-purification, and printing. The democratization of kinetic profiling will be facilitated through the provision of SPOC protein chips to both research and industry users as catalog array and off-the-shelf protein chips, expediting their proteomics kinetics research workflow. Further, we have expanded our capacity for flexible sensor development by creating and applying self-assembled monolayers on bare gold slides which allows us to produce gold sensor chips with custom surface chemistries and to tailor SPOC technology according to user needs and applications. We have developed processes to fabricate gold SPRi slides from scratch to help reduce the overall cost of technology development and the SPR chips offered to customers in future. The SPOC platform will enable researchers to customize protein panels for HTP applications such as monoclonal antibody characterization, AI-designed drug validation, and preclinical drug development to screen libraries of drug candidates against thousands of on-and off-protein targets. Other potential application areas include determination of basic protein-protein interaction kinetics, vaccine efficacy monitoring, serum profiling, biomarker discovery, and diagnostics.

### Methods

#### 1. DNA preparation, SARS-CoV-2 RBD mutants, Recombinant plasmids construction

Genes of interest were cloned into *in vitro* transcription and translation-compatible expression vectors, including pJFT7_nHALO (or cHALO) from DNASU plasmid repository, Biodesign Institute, Arizona State University and custom designed pT7CFE1_nHalo (or cHalo) vectors. These expression vectors ensure *in vitro* expression of proteins of interest with an N-or C-terminal HaloTag for immobilization on HaloTag chloro-alkane linker functionalized surfaces. Most of the recombinant plasmids expressing the genes of interest were either procured from DNASU plasmid repository or outsourced to our commercial vendors – Genscript and Twist Biosciences for custom syntheses. Dispense plates for DNA printing were prepared with recombinant plasmids at a concentration of 100 ng/µL.

#### 2. Preparation of silicon nanowell substrate for DNA printing

Silicon slides consisting of thousands of nanowells were fabricated at a semiconductor foundry and prepared for DNA printing by vapor-phase deposition of (3-Aminopropyl)-triethoxysilane (APTES) on the surface using an internally optimized protocol. To clean the silicon nanowell slides prior to printing, slides were placed in 50 mL conical tubes containing distilled water and then centrifuged at 1,400 *x g* for 2 minutes, inverted and repeated (Premiere, Model XC-2450 Series Centrifuge). Afterwards, slides were dried with a nitrogen gun and then subjected to oxygen plasma treatment at 50W for 2 mins using a Plasma Etch instrument (Plasma Etch Inc., Model PE-100HF). The nanowell substrates were immediately placed in a vacuum desiccator containing 2 mL APTES in a vial, subjected to vacuum/filling for a total of 15 min using low pressure nitrogen, and eventually incubated for 1.5 h under vacuum. Afterwards, APTES-modified nanowell substrates were removed, cured at 100°C for 1 h, and stored in a Bel-Art autodesiccator (catalog number 420740117).

#### 3. Immobilization of unique expression plasmids in nanowells

DNA printing was outsourced to Engineering Arts, LLC and performed on the RM3 system. The technique employs high-speed piezo printing with an au302 piezo dispense system, featuring an integrated alignment system designed for microwell, micrometer angular alignment fixture, look down camera, transfer arm, and a vacuum tray (Karthikeyan et al., 2016). First, silicon nanowell slides were placed and aligned on the vacuum tray using the robotic transfer arm, micrometer angular alignment fixture, and look down camera. Print mix was then printed in picoliter quantities “on-the-fly” using a multi-dispense head into each nanowell, followed immediately by printing of picoliter quantities of expression plasmid DNAs from the dispense plates. The print mix contained HaloTag chloro-alkane linker (O4) amine, which was varied from 0 to 1 mM (Promega, P6741), 1.46 mg/ml BS3 in DMSO, and 7.4 mg/mL BSA, in nuclease-free water. The addition of HaloTag chloro-alkane linker into the print mix was used for limited capture of expressed HaloTagged proteins at the bottom of nanowells for expression validation using anti-HaloTag antibody via immunofluorescence. After the piezo printing process, the nanowell slides were stored in an autodesiccator until they were ready for use.

#### 4. Bio-functionalization of SPOC slides with HaloTag chloro-alkane linker

Bare gold-coated slides compatible with SPRi biosensing were purchased commercially, in addition to hydrogel coated slides purchased from Xantec (HC30M). Bare gold slides used in OpenPleX SPOC array analysis were derivatized with a self-assembled monolayer (SAM) using a 2 kDa Thiol-PEG-NHS reagent (BioPharma PEG; HE003023-2K) towards HaloTag chloro-alkane linker modification and Halo-protein capture. For SAM assembly, 20 μl of 200 μM Thiol-PEG-NHS solution in DMSO was pipetted onto a clean microscope slide and the bare gold SPR slide was placed gold-side down onto the solution. After 2 h incubation at room-temperature, the slide was rinsed briefly in 200 Proof Ethanol and distilled water and dried with compressed air. For preparation of HC30M slides for Carterra LSA^XT^ SPOC array analysis, sensors were activated with a 1:1:1 mixture of 0.4M N-(3-dimethylaminopropyl)-N-ethylcarbodiimide (EDC), 0.1 M N-hydroxysuccinimide (NHS), and 0.1M 2-(N-morpholino) ethanesulfonic acid, pH = 5.5 (MES) for 10 minutes at room-temperature. For either Thiol-PEG-NHS derivatized sensors or HC30M hydrogel slides, after rinsing with distilled water and drying, the slides were incubated for either 1 hour or overnight with 1 mg/mL of HaloTag chloro-alkane linker (Iris Biotech; RL-3680). The sensors were then incubated for 30 min 0.5 M Ethanolamine (pH 8.5) to quench any remaining free-NHS esters.

#### 5. Automated in-situ production of SPOC array

SPOC arrays were produced via automated IPC using a custom AutoCap instrument designed and built in-house. Nanowells printed with DNA plasmids of interest were filled via centrifugation in 50 mL conical tubes containing water for 4 min on each end at 1,400 x *g*, followed by incubation in Superblock (Thermofisher Scientific) for 30 min with gentle rocking. SPOC arrays were blocked for 30 min in 0.5 M ethanolamine, pH 8.5 to quench any free ester groups. After blocking, both the nanowell and SPOC slides were rinsed at least 3 times with water and dried gently with compressed air. 1-Step Human Coupled IVT mix was prepared and centrifuged for 20 min at 19,000 RCF at 4°C to remove particulates. The supernatant was gently removed to a fresh tube and stored at 4°C until injection into the instrument. The automated IPC instrument contains four stations, each one capable of holding one 25 mm x 75 mm nanowell slide. Each station contains a hydraulic actuator-driven base that holds a nanowell slide surrounded by a silicone gasket. A hinged lid designed to fit the SPOC arrays in a precise location above the nanowell slide is fitted onto the base of each station. Automated software was designed in-house using LabView to control all mechanical parts of the instrument. For protein expression and capture, the nanowell slide was placed on the base and the SPOC slide was mounted to the lid. The lid was then affixed firmly to the base using wingnuts and, using the automated software, low pressure (20 psi) was applied via the actuator, creating an airtight chamber containing the two slides with the SPOC array positioned immediately above the nanowell slide with a small gap in between. Following application of low pressure, the chamber and fluidic lines were vacuumed for a total of 12 min, with the goal of evacuating air from the nanowells and chamber. Using a primed 500 µL Hamilton syringe, the lysate was aspirated into the chamber via vacuum following automated valve switching from vacuum line to the injection line. This process results in the nanowells and the small gap between the slides being immediately filled with the lysate mixture. Once the slide space is filled (less than 10 sec), high pressure (220 psi) is applied via the actuator to tightly press-seal the two slides together. The instrument space was closed and brought to 30°C for 2-4 h for *in vitro* translation and expression (IVTT), with the nanowells serving as nanoreactors for IVTT and the resulting HaloTagged protein being concurrently captured onto the SPOC array. Following IVTT, the pressure is removed, and the slides are immediately washed in cold 1X PBST 3 times prior to assay to remove any unbound IVTT components and incompletely expressed protein.

#### 6. Protein analysis on surface plasmon resonance instruments

SPR experiments were performed on Carterra LSA^XT^ SPR and Horiba Scientific OpenPleX SPRi instruments. SPOC protein array chips from the expression/capture process were washed with 1X PBST and deionized water and dried with compressed air prior to prism mounting and instrument installation.

##### 6.1 Sensor response assay protocol using Carterra LSA^XT^

The sensor for SPOC Carterra LSA^XT^ analysis was prepared by mounting the SPOC chip onto a custom research-use-only plain-prism Carterra LSA^XT^ cartridge using 15.0 µL of a refractive index matching oil (Cargille. Cat# 19586) and then inserted into the instrument. Once installed, the single flow cell was docked onto the sensor and the sensor was primed with running buffer (1X PBS pH 7.3, 0.2% BSA, 0.05% Tween-20; filtered and degassed). Sensor temperature was maintained at 15°C for the duration of the experiment. Regions of interest (ROI) were manually assigned over spots of interest in reference to the spot array orientation mark (hockey stick mark). Antibody titrations were performed by diluting antibodies into running buffer followed by performing 3, 3-fold serial dilutions resulting in a total of 4 serially diluted analyte samples with 300 µL volumes each. Association and dissociation rates varied from 4 to 30 min and 8 to 45 min, respectively and all screens were performed using the QuickStart experiment menu of the Carterra LSA^XT^ control software.

##### 6.2 Sensor response assay protocol using the OpenPleX SPRi

The SPOC chip was mounted onto a Horiba glass prism using 0.5 µL of high index oil (Serie B, Part# 1300007816, Horiba Scientific). The sensor-coupled prism was installed into the instrument by manually pressing onto a gasketed flowcell. The SPR Running Buffer for OpenPleX consisted of freshly prepared 1X PBS added with 0.2% BSA, 0.05% Tween-20, filtered using a 0.22 µm bottletop filter (Corning; CLS430626) and de-gassed for 20 min. SPR Running Buffer was initially flushed over the SPOC protein chip at a high flow-rate of up to 2,000 µL/min to wet the sensor surface and remove bubbles. Afterwards, the buffer flow rate was reduced to 50 µL/min and the flowcell temperature was set at 25°C for the remainder of the experiment. After ROI selection and instrument calibration with 1% glycerol in the running buffer, 200 µL analyte in running buffer was flowed over the sensor surface via a sample injection loop with the association phase performed for 3.5 min followed by 5 min dissociation phase for all blank and analyte injections. Protein expression/capture on the SPOC chip was confirmed by flowing 200 µL of mouse anti-HaloTag (133 nM) in running buffer. Binding to protein-specific antibodies and cross reactivity was validated by injecting mouse anti-Jun (200 nM) and mouse anti-p53 (17.8 nM).

*<u>On-slide post translational citrullination</u>* of four spots of unmodified Halo-HIST1H3A proteins on SPOC array was performed on the OpenPleX instrument. Prior to this reaction, the chip which also contained two protein spots each of Halo-RBD BA.1 and Halo-CT45A3 as controls was probed with anti-Halo and 200 µL of 1:50 rabbit anti-citrullinated HIST1H3A in running buffer to confirm protein expression and the absence of citrulline group on the native HIST1H3A proteins. The citrullination reaction was performed on the chip while mounted on OpenPleX instrument by re-circulating for 2 h at RT a 1 mL citrullination reaction mixture containing 0.3 U/mL Peptidyl Arginine Deiminase (Sigma P1584-25UN), 10 mM CaCl2, 100 mM Tris pH 7.6, and 5 mM DTT. After buffer wash, the anti-cit-HIST1H3A was injected, followed by secondary anti-rabbit antibody injection to detect citrullinated HIST1H3A. The difference image was captured and graph of reflectivity change for the HIST1H3A, BA.1 RBD and CT45A3 spots was plotted.

##### 6.3 Extraction of HTP Kinetic Information

Raw data from the Carterra LSA^XT^ were analyzed in Kinetics analysis software (Carterra, v1.9.0.4167), while data from the OpenPleX were processed using ScrubberGen2 (BioLogic Software, v2.0g). In all cases, the data were y-aligned and double-referenced via blank, running buffer alone injection subtraction and referencing of the data against control spots of the array where no binding was expected and observed. Once pre-processed, all data were globally fit in the software using a 1:1 Langmuir binding model to obtain kinetic parameters and equilibrium dissociation constants. Since the analytes used in the assay are bivalent, the data is complicated by avidity effects and therefore the 1:1 binding model is acknowledged to yield only an approximation of the affinity for each antibody-antigen interaction interrogated.

##### 6.4 Regeneration of SPOC Chips

Different sensor regeneration conditions and buffers were tested to validate functionality of SPOC proteins, including 0.5-2 M NaCl in 5 mM EDTA, 10-100 mM HCl, and 10 mM glycine-HCl, pH 2.4. In this experiment, SPOC array of 132 HaloTagged protein spots was mounted on Carterra LSA^XT^, followed by injection of anti-HaloTag antibody. The sensor surface was regenerated using the different buffers and then probed again with anti-HaloTag. The process was repeated three times. In each case, average response units ± standard deviation from all 132 Halo-protein spots on the array was calculated. A bar graph of the average response units for the initial anti-Halo injection and all three regenerations was plotted (**Supplemental Figure S1**).

#### 7. Determination of differential binding kinetics of a commercial anti-SARS-CoV-2 Receptor Binding Domain (RBD) antibody against SARS-CoV-2 RBD using SPOC

Recombinant plasmid DNAs encoding SARS-CoV-2 RBD domain variants Alpha, Delta, Omicron variants and subvariants BA.1, BA.5, BA.2.75, BQ.1, XBB.1.16, XBB.1.5 and BQ.1.1, and the wildtype Wuhan RBD as HaloTag genes were printed in silicon nanowell slides. Using the SPOC protein array production platform, RBD variants were expressed *in situ* and capture-purified onto replicate SPOC chips as Halo-fusion protein arrays. The SPOC chip was screened on the Carterra LSA^XT^ SPR instrument. Protein expression/capture was validated by injecting 133 nM of anti-HaloTag antibody, followed by surface regeneration using 2 M NaCl and buffer wash prior to probing with anti-RBD antibody. SPOC RBD chips were screened with two different manufacturer lots of a commercial mouse anti-RBD from Proteintech Group (catalog number: 67758-1) to determine the differential binding kinetics to the RBD variants. To ensure reproducibility, assays were performed multiple times using SPOC chips produced at separate protein expression/capture runs and screened weeks apart. Also, serial concentrations 19.75, 59.26, 177.78, and 533.33 nM of each anti-RBD Lot was injected in succession to confirm sensor response curves. In the first replicate SPOC assay, SPOC chip containing the Wuhan wildtype RBD and five RBD variants – Alpha, Delta, Omicron BA.1, BA.5, BA2.75 was probed with the mouse anti-RBD from manufacturer Lot 1. The second and third assays were similar and performed with SPOC chips containing all nine variants and the Wuhan wildtype and probed with the mouse anti-RBD from Lot 2. The association/on-rate (10 min) and dissociation/off-rates (15 min) kinetics were monitored during injections, and sensor surface was regenerated using 2 M NaCl between injections. The differential binding was validated orthogonally by immunofluorescence assay using the mouse anti-RBD and secondary goat anti-mouse Cy3.

#### 8. Antibodies used for label-free and fluorescent SPOC assays

Mouse anti-HaloTag antibody utilized for confirming expression of IVTT expressed HaloTag-fusion proteins by SPR was purchased in glycerol-free format from Chromotek (28a8), while a rabbit anti-HaloTag from Promega was used for fluorescent-based expression validation (G9281). Protein specific antibodies used in this study include mouse anti-Jun (ThermoFisher, 39-7500), mouse anti-p53 (Sigma-Aldrich, P6874), mouse anti-Src (ThermoFisher, AHO1152), mouse anti-RBD (Proteintech, 67758-1), rabbit anti-Fos (14C10) rabbit anti-citHIST1H3A (Abcam, ab5103), rabbit anti-citFGA (ImmunoPrecise, MQ13.102), and anti-phosphoTyrosine (ThermoFisher, 03-7700). For fluorescent assays, secondary antibodies used include goat anti-Rabbit-Cy3 (Jackson ImmunoResearch, 111-165-003) and goat anti-Mouse-Cy3 (Jackson ImmunoResearch, 115-165-062).

#### 9. Fluorescence assays to validate functionality of SPOC proteins captured on regular 25 mm x 75 mm glass slide

*<u>Validation of Halo-protein capture and binding to protein-specific antibodies</u>* on SPOC glass slide was confirmed via immunofluorescence by incubating the whole slide in 5% milk in 1X PBST containing 1:750 rabbit anti-HaloTag or protein-specific antibodies (anti-p53, Jun, or Fos). After 30 min – 1 h of incubation at RT and washing with PBST, the slide was again incubated for 30 min with 1:500 dilution of the appropriate secondary antibody fluor-conjugate in 5% milk-PBST. After the secondary incubation, the slide was washed with PBST, rinsed with water, dried with compressed air, and scanned using an InnoScan 910 AL Microarray Scanner (Innopsys, Carbonne, France). *<u>In vitro citrullination of SPOC proteins</u>*was performed by incubating SPOC glass slides containing Halo-FGA and Halo-HIST1H3A with the citrullination reaction mixture containing 0.3 U/ml Peptidyl Arginine Deiminase (Sigma P1584-25UN), 10 mM CaCl2, 100 mM Tris pH 7.6, and 5 mM DTT. After incubation for 2 h and buffer wash, citrullinated FGA and HIST1H3A on the SPOC glass slide were probed by incubating for 1 h with 1:500 rabbit anti-citrullinated HIST1H3A or FGA and detected with goat anti-rabbit Cy3 using the InnoScan 910 AL Microarray Scanner. PAD2 was omitted from control slides. *<u>Protein-Protein Interaction assay on glass</u> <u>slides</u>* were performed by probing a whole SPOC glass slide containing two Halo-Jun and six Halo-Fos protein spots with buffer solution containing 10 ug/mL commercial recombinant Fos protein in 1% milk-PBST for 1 h at RT. Binding of the recombinant Fos protein to the SPOC Jun was detected by incubating the SPOC glass slide with 1:500 rabbit anti-Fos antibody in 5% milk PBST and then goat anti rabbit Cy3, followed by fluorescence scanning with the InnoScan 910 AL. *<u>Enzymatic activity assays on SPOC glass</u> <u>slides</u>* were performed using dephosphorylation and autophosphorylation of Src kinase on SPOC array. To ensure that endogenous phosphorylation of Src from IVTT expression is not being detected, the slide was first dephosphorylated by incubating for 30 min at 37°C with assay mixture containing 0.5 units of calf intestinal alkaline phosphatase in 1X dephosphorylation buffer (Thermo Scientific, 18009019) and 1 mM MgCl2, followed by 3 washes with Tris-buffered saline. The removal of phospho groups from immobilized Src kinase was confirmed by staining with anti-phosphotyrosine antibody. Next, auto-phosphorylation of the Src was performed by incubating the SPOC glass slide for 45 min at RT in phosphorylation buffer containing 4 mM ATP, 50 mM Tris pH 7.6, 5 mM MgCl2, and 0.5 mM DTT. The phosphorylated Src was then detected via anti-phosphotyrosine antibody.

## Acknowledgement

This work was supported by the National Institute of Health SBIR grants 1R43OD024970-01A1 and 1R44TR004297-01, and SPOC Proteomics’ internal funding. We are immensely grateful to Dr. Hwall Min, whose invaluable contributions were pivotal in the initial validation of the SPOC technology. Our gratitude extends to the Carterra R&D team, particularly Dr. Rebecca Rich and Noah Ditto who supported the integration of SPOC proteomic biosensor platform with the Carterra LSA^XT^ instrument.

## Author Contributions

B.T conceived the experiments. W.M, C.V.A, and R.C performed the experiments, analyzed the data and prepared the figures. M.M, L.G helped with project planning and joined the discussions. C.V.A drafted the manuscript. All authors reviewed and approved the manuscript.

## Data availability statement

The authors confirm that the data supporting the findings of this study are available within the article and its supplementary materials.

## Notes

### Competing Interest Statement

The authors have declared no competing interest.

