## Supplemental Figures for "Novel sensor-integrated proteome on chip (SPOC) platform with thousands of folded proteins on a 1.5 sq-cm biosensor chip to enable high-throughput real-time label-free screening for kinetic analysis"

**SUPPLEMENTAL DATA**

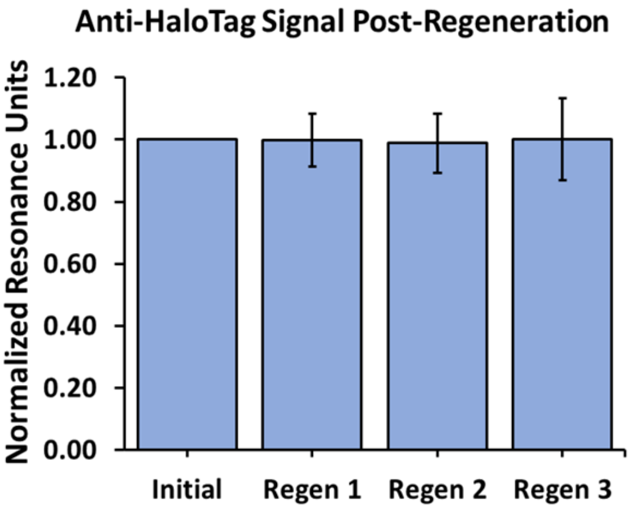


**Figure S1 | Reproducibility of SPOC sensor binding signal post-regeneration.** The bar plot shows the normalized Mouse anti-HaloTag Rmax signal observed after four injections of the antibody measured for each of 132 spots across the SPOC array (± SD). The sensor was regenerated with 10mM Glycine-HCl (pH = 2.4) between each injection and greater than 99% of the HaloTag signal was retained between regenerations for all ligand containing spots. The regeneration experiment was performed using the Carterra LSA$^{XT}$ SPR instrument.

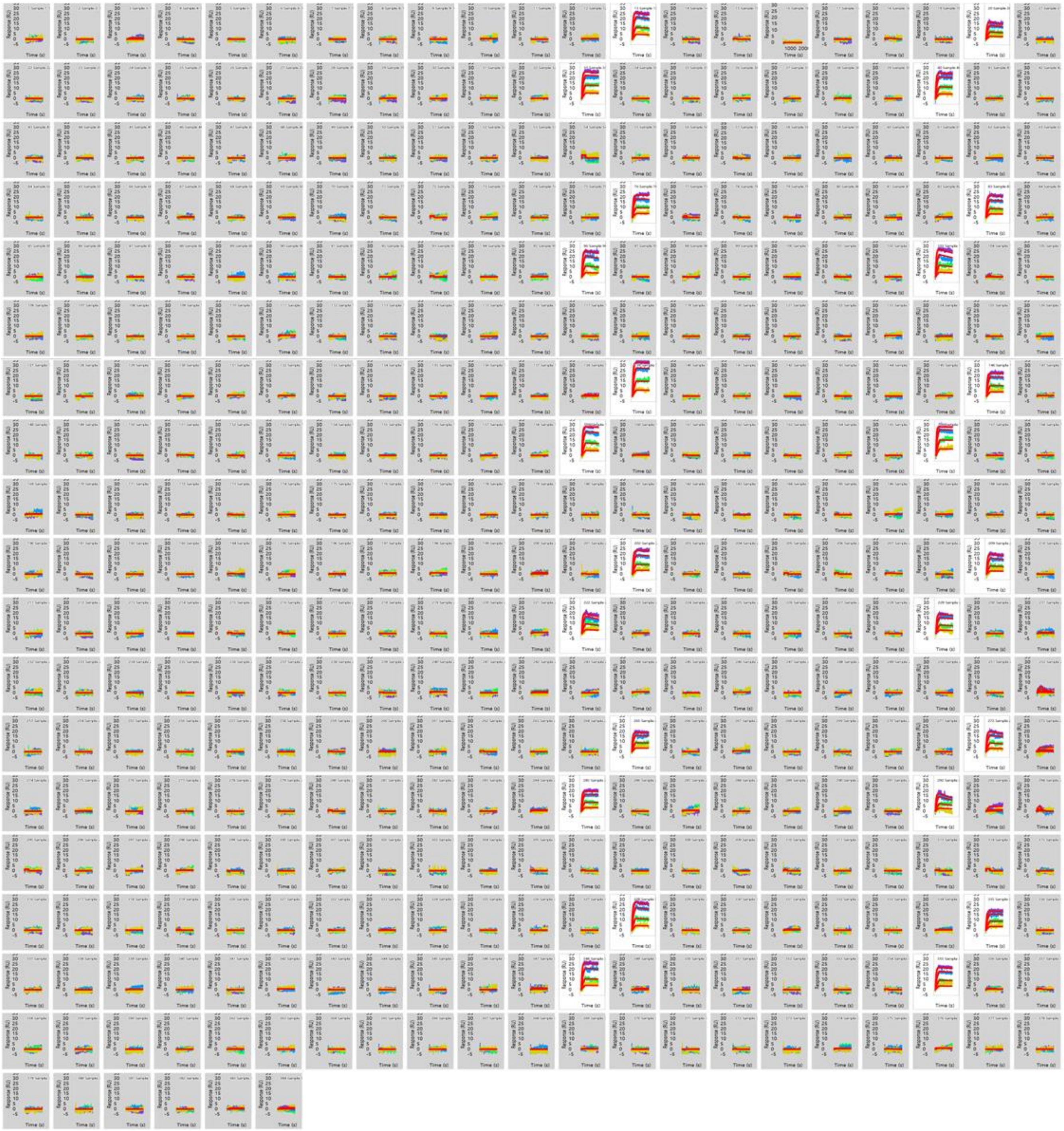

**Figure S2 | Enlarged view of all Carterra LSA$^{XT}$ sensorgrams from 384 spots across the 30k array after injection of mouse anti-p53 antibody titrations.** Only the twenty-four replicate spots where p53-HaloTag was expressed and captured responded to the anti-p53 injections (un-shaded sensorgrams).

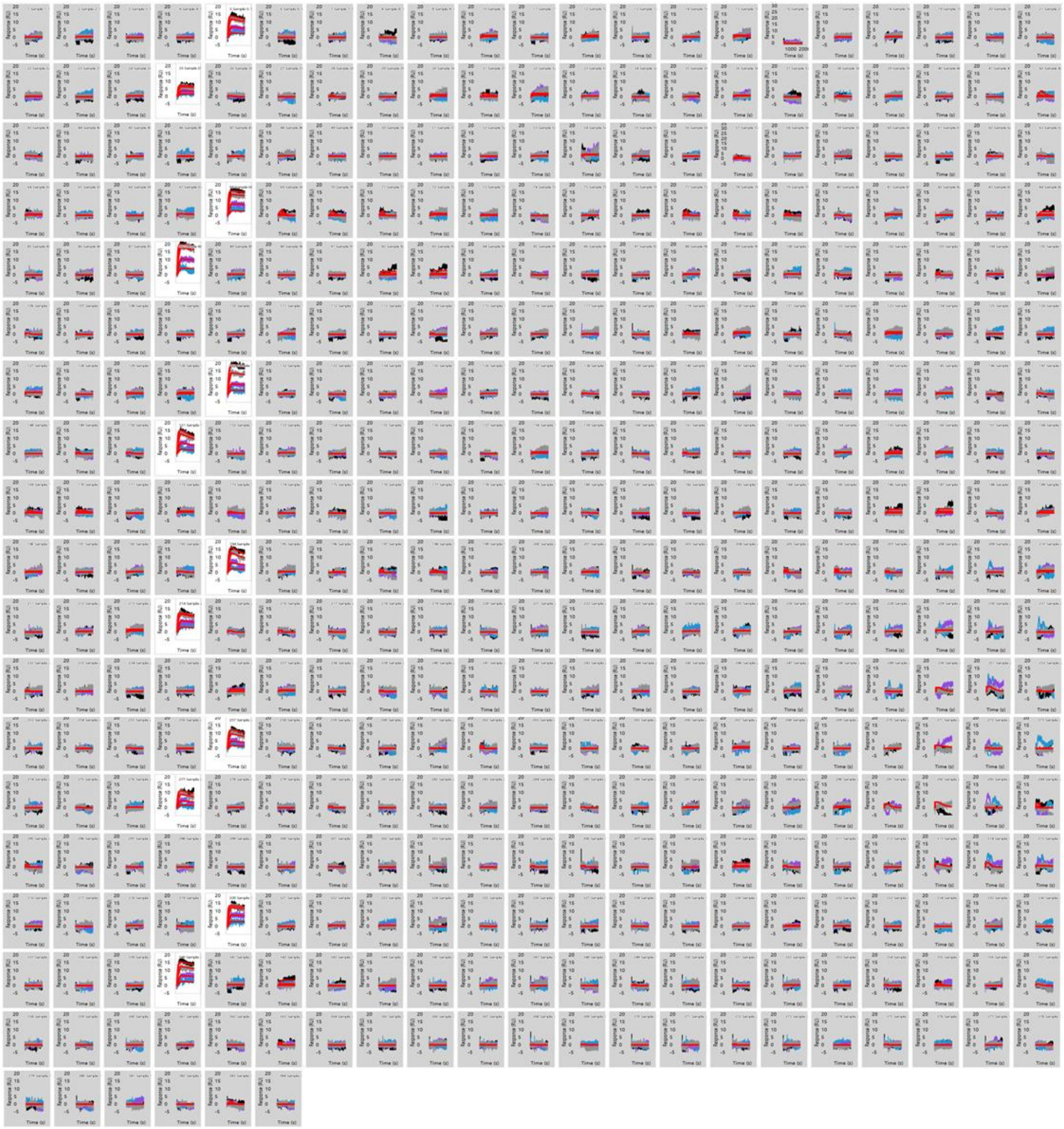

**Figure S3 | Enlarged view of all Carterra LSA$^{XT}$ sensorgrams from 384 spots across the 30k array after injection of mouse anti-Src antibody titrations.** Only the twenty-four replicate spots where p53-HaloTag was expressed and captured responded to the anti-p53 injections (un-shaded sensorgrams).

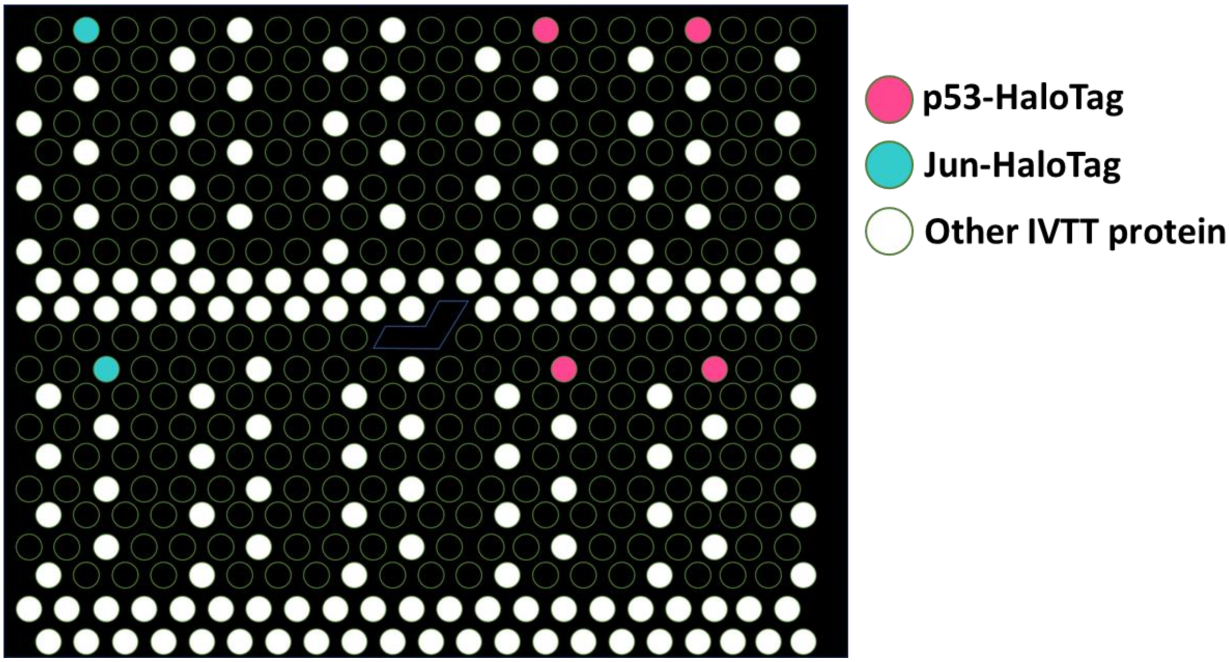


**Figure S4 | Spot map for the SPOC array analyzed on the OpenPleX SPRi instrument.** The print locations for the IVTT compatible plasmid DNA encoding HaloTag fusion proteins are shown in white and were printed in the nanowell slide as 2 replicate subarrays above and below the reference mark with each sub-array printed in a sparse-dense pattern. This print contained four total spots for expression of p53-HaloTag (pink) and two spots for expression of Jun-HaloTag (blue).

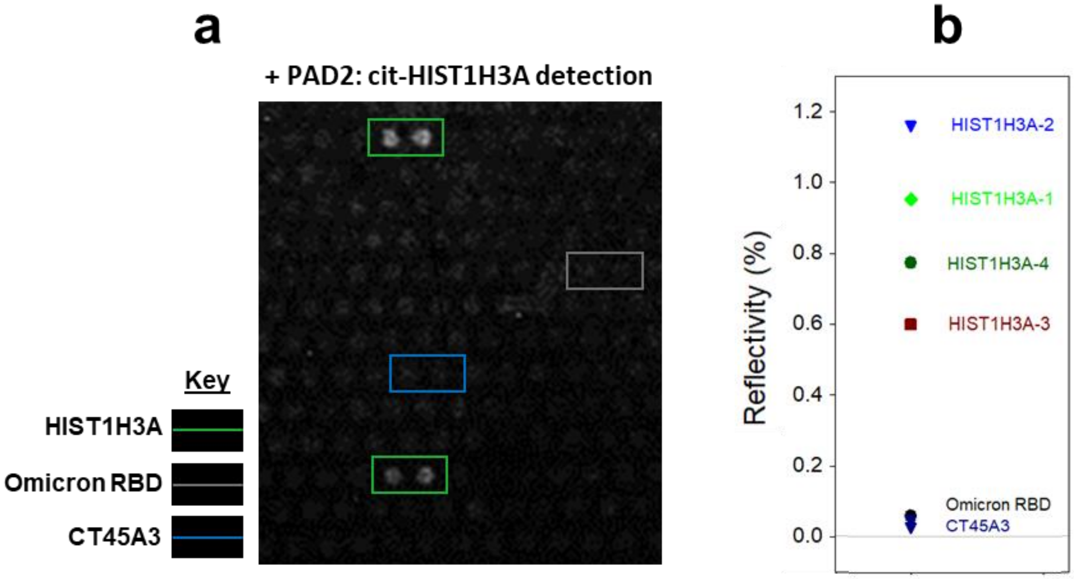


**Figure S5 | OpenPleX SPRi validation of *in vitro* PAD2 citrullinated SPOC proteins.** (**a**) Difference image shows detection of cit-HIST1H3A after the arrayed biosensor was treated with PAD2 for 1 hour at room-temperature while installed in the OpenPleX SPRi instrument. (**b**) Graph showing reflectivity change measured for each protein species in a after on-slide PAD2 treatment and injections of anti-cit-HIST1H3A and secondary antibodies. All 4 HIST1H3A spots are disproportionately detected, indicating that citrullination of proteins captured on biosensor surfaces is feasible as on glass slides.

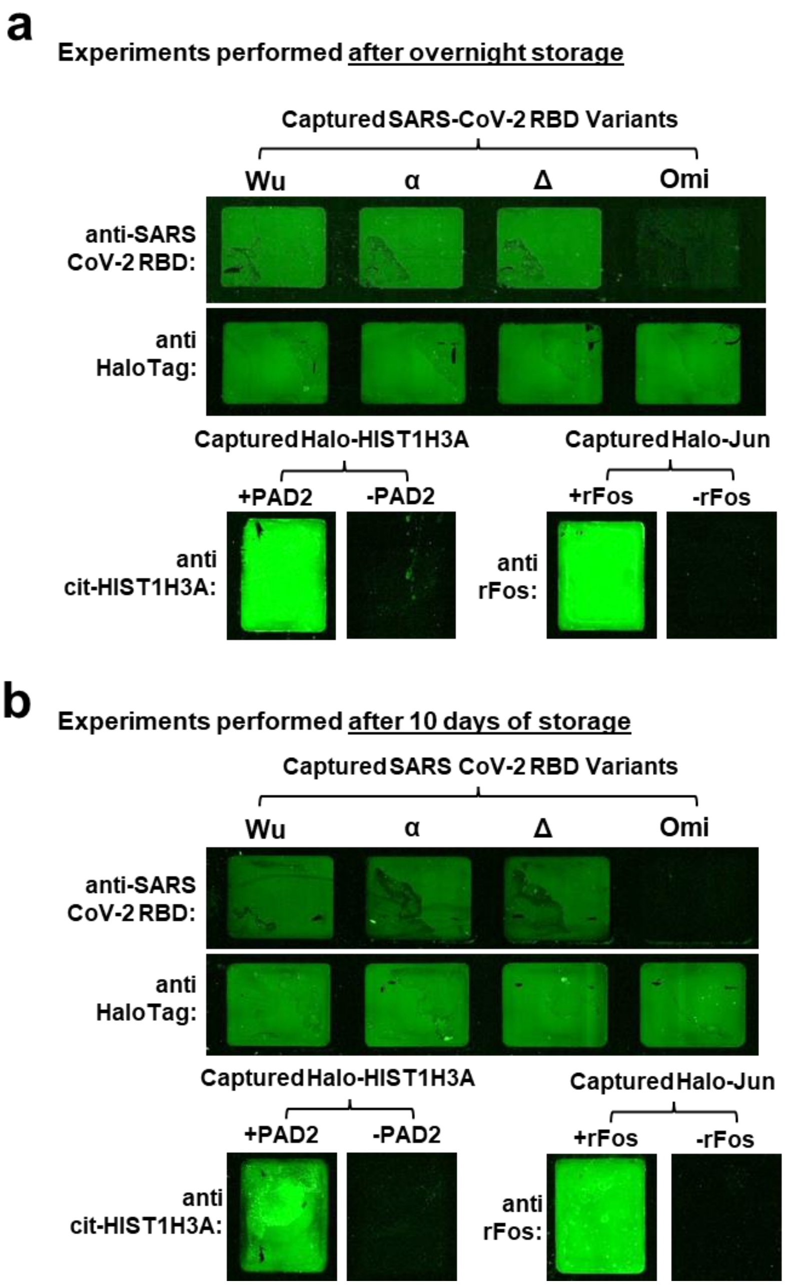


**Figure S6 | Fluorescence assay on glass surfaces demonstrating IVTT expressed and capture-purified proteins are stable and active for 10 days in storage.** IVTT expressed HaloTag fusion proteins were manually captured onto a functionalized glass slide and a variety of assays were performed including (i) detection of CoV-2 RBD variants with primary and secondary antibodies, (ii) PAD2 mediated citrullination of IVTT expressed HIST1H3A, and (**iii**) capacity for recombinant Fos protein (rFos) to interact with IVTT expressed Jun. Fluorescent secondary antibodies were used to detect all bound proteins and their modification. Slides with captured IVTT proteins were prepared and stored (**a**) overnight at 4°C in PBS supplemented with 5% glycerol before assay or (**b**) stored for 10 days at -20°C in PBS with 50% glycerol prior to assay.

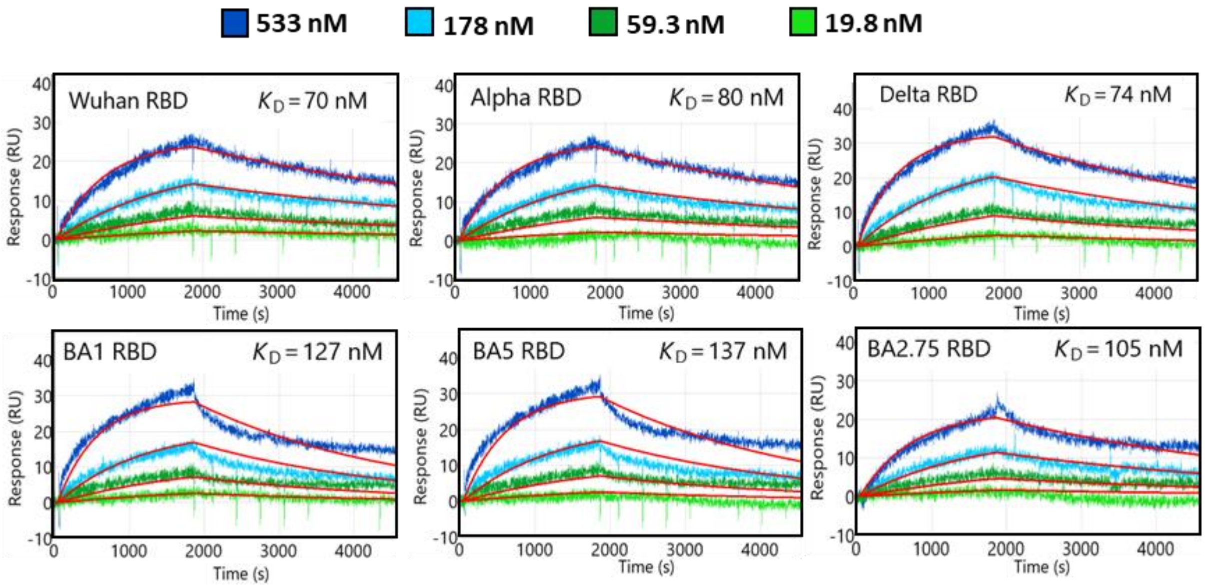


**Figure S7 | Carterra LSA$^{XT}$ Kinetic Data of Proteintech Mouse anti-RBD from Lot 1.** Sensorgrams from the mouse anti-RBD injections against 6 RBD VoC recorded from a SPOC array are shown with 1:1 binding model overlayed (red curved) and equilibrium dissociation constants displayed above each respective sensorgram.
